# Recurrent sex chromosome turn-over in termites

**DOI:** 10.1101/2024.08.12.607539

**Authors:** Roxanne Fraser, Ruth Moraa, Annika Djolai, Nils Meisenheimer, Sophie Laube, Beatriz Vicoso, Ann Kathrin Huylmans

**Author notes:** These authors contributed equally to the manuscript (Roxanne Fraser, Ruth Moraa).

## Abstract

Termites, together with cockroaches, belong to the Blattodea. They possess an XX/XY sex determination system which has evolved from an XX/X0 system present in other Blattodean species such as cockroaches and wood roaches. Little is currently known about the sex chromosomes of termites, their gene content, or their evolution. We here investigate the X chromosome of multiple termite species and compare them to the X chromosome of cockroaches using genomic and transcriptomic data. We find that the X chromosome of the termite *Macrotermes natalensis* is large and differentiated showing hall marks of sex chromosome evolution such as dosage compensation, while this does not seem to be the case in the other two termite species investigated here where sex chromosomes are probably evolutionary young. Furthermore, that X chromosome in *M. natalensis* is different from the X chromosome found in the cockroach *Blattella germanica* indicating at least one, potentially multiple, sex chromosome turn-over events during termite evolution.

## Introduction

The mode of sex determination is one key difference between termites and the well-studied other big group of social insects, namely Hymenoptera to which ants and bees belong. In the termites, males are usually present in all castes (1), do not solely serve reproductive purposes (2), and are diploid like females (3, 4). Sex determination has been shown to be based on a male-heterogametic XX/XY system which could be documented for a wide number of termite species (5–7). Phylogenetically, termites (*Isoptera*) form a monophyletic group nested in the paraphyletic cockroaches (*Blattaria*) together forming the order Blattodea (4, 8, 9). Cockroaches differ from termites in their sex determination system and have been shown to possess an XX/X0 system where the male possesses only one X chromosome but no Y chromosome (10, 11). This leads to the question how the XY system in termites has evolved. As the XX/X0 system could be demonstrated for multiple cockroach species (summarised in (7)), it is unlikely that the Y chromosome was lost in all these species independently. The wood roaches of the genus *Cryptocercus* are the sister group to the termites. Their sex determination system has also been described as an XX/X0 system as demonstrated by karyotyping data (12–14). Hence, it can be assumed that the Y chromosome appeared after *Isoptera* split from *Cryptocercoidae* (star in Fig. 1 A). In consequence, XX/X0 was probably the ancestral state and an event during the evolution of the termites must have led to the emergence of the Y chromosome approximately 150 million years ago (9, 15, 16).

**Fig. 1.**
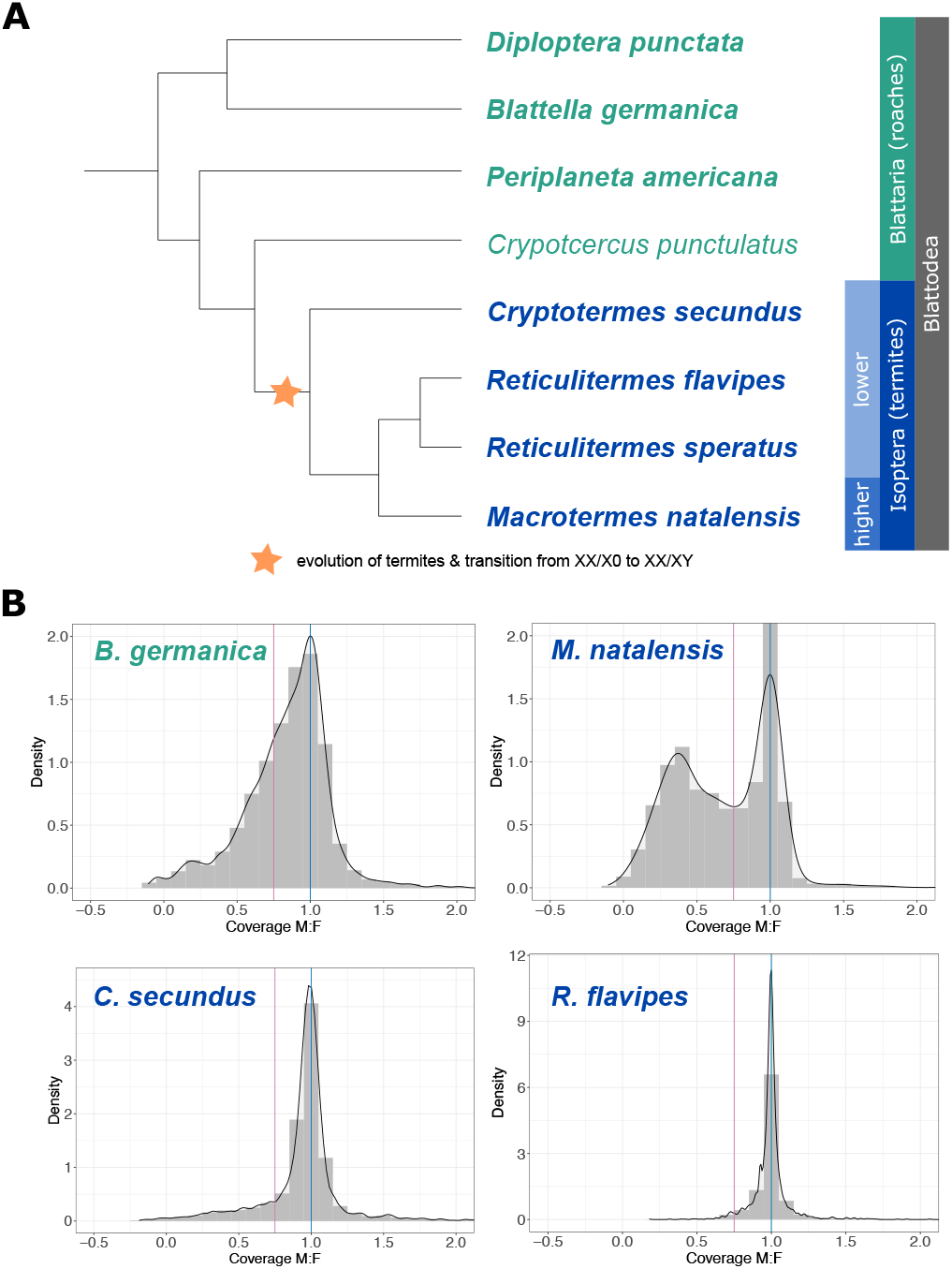
Sex chromosome evolution in Blattodea. (A) Phylogenetic tree showing relationships between selected wood roach, cockroach, and termite species. Note that while the tree includes wood roaches (represented by *Crypotocercus punctulatus*) to illustrate their relationship with the other species, due to data availability only cockroach and termite species were included in the analysis (species names in bold). Paraphyletic roaches are represented in green, social termites in blue. The star marks the assumed transition between XX/X0 sex determination in roaches and XX/XY in termites. (B) Genomic coverage per scaffold plotted as male-to-female ratio for the cockroach *B. germanica* and the three termite species *M. natalensis, C. secundus*, and *R. flavipes*. Distributions were centered with the autosomal peak (blue vertical line) at one and the cut-off for the X chromosome identification is shown (pink vertical line).

Most of the data on sex determination systems in Blattodea are based on karyotyping or cytogenetic analyses that yield information about DNA content differences, chromosome morphology, and numbers differing between the sexes (reviewed and summarised in the Blattodea kayrotype database (7)), but to date, little information on the genomic level such as gene content, genetic differentiation, or gene expression is available for this group. Only in cockroaches have some studies looked at the X chromosome from a genomic perspective (e.g. (17, 18)). However, data was (and often still is) limited for these species making conclusive inferences about processes such as dosage compensation in the German cockroach *Blattella germanica* complicated (17). In termites, amongst the six published genomes, females were chosen for sequencing in three cases (*Reticulitermes speratus* (19), *Reticulitermes lucifugus* (20), and *Macrotermes natalensis* (21)). Genomes for *Cryptotermes secundus* (4), *Zootermopsis nevadensis* (22), and *Coptotermes formosanus* (23), on the other hand, were based on genomic DNA extracted from many pooled individuals including males. Hence, we expect Y-derived scaffolds to only be present in these three genome assemblies. While some studies on gene expression differences between males and females exist (19, 22), current genome studies have paid little attention to the sex chromosomes. Thus some interesting questions about their evolution remain unanswered.

In insects, different to other groups such as mammals or birds, it has been shown that sex determination systems can be very variable and frequently turn over (24, 25). The mechanisms through which this occurs are manifold and are often hard to observe in action (reviewed in e.g. (26)). In termites, the Y chromosome suddenly appears that presumably did not exist in the ancestor with cockroaches and wood roaches. For the transition of an XX/X0 system to an XX/XY system, multiple different scenarios are possible (analogous to the ZZ/Z0 to ZZ/ZW transition in Lepidoptera (27)): there could have been a (i) complete turn-over of sex chromosomes at the base of *Isoptera* making a pair of formerly autosomal chromosomes the new X and Y chromosomes, (ii) chromosomal fusion of the ancestral X chromosome to an autosome making the homologous chromosome a neo-Y chromosome, or (iii) recruitment of an extra-genomic element such as a B chromosome.

These three different scenarios lead to different predictions regarding the conservation of X-linked sequences from cockroaches to termites. Under scenario (i), we expect cockroach X-linked genes to be autosomal in termites and vice versa. Under scenario (ii), we expect X-linked genes to still be X-linked in termites but we expect to find additional genes to be X-linked in termites that were previously autosomal in cockroaches, preferentially coming from a single cockroach chromosome. Finally, under scenario (iii), we expect the X chromosomes of cockroaches and termites to be largely conserved in size and gene content.

We investigate here these possible scenarios for the origin of the Y chromosome using the X chromosome of termites as currently no chromosome-level genome assemblies of termites are available and current genomes often lack Y-chromosomal sequences. For this, we use a coverage approach where we sequenced DNA from males and females of three different termite species in order to identify the X chromosome and compare them to the cockroaches. We also assess gene expression in both sexes to study dosage compensation of the X chromosomes. We find that only one termite species clearly shows a differentiated large X chromosome that, however, is different from the X chromosome of the German cockroach indicating that turn-over of sex chromosomes happened in termites.

## Methods

### Sequencing

Reproductive individuals from lab colonies of three termite species, *Reticulitermes flavipes, Macrotermes natalensis*, and *Cryptotermes secundus*, were supplied and sexed externally by our collaborators Dino McMahon, Mireille Vasseur-Cognet, and Judith Korb, respectively. Subsequently, DNA of a male and a female each was extracted with the Qiagen DNeasy Blood & Tissue kit. Library preparation and paired-end 125 base-pair (bp) whole-genome DNA-sequencing was performed on an Illumina HiSeqV4 at Vienna Biocenter Next Generation Sequencing Core Facility. All raw DNAseq data are publicly available in the NCBI Short Read Archive (SRA) under the BioProjects **XXX**.

### Sex scaffold classification

DNA reads were mapped to their respective references genomes using Bowtie2 (version 2.3.5.1; (28)). The cockroach reference genomes are available at NCBI under GCA_003018175.1 (version 1) for *B. germanica* (4), GCA_025594305.2 (ASM2559430v2; version 2) for *P. americana* (29), and GCA_030220185.1 (version 1) for *D. punctata* (30). The reference genome of the termite *C. secundus* is available at NCBI under GCF_002891405.2 (version 2) (4), and that for *M. natalensis* can be downloaded from GigaScienceDB under https://doi.org/10.5524/100055 (21). Since no reference genome is currently available for *R. flavipes*, genomic reads were mapped to the assembled genome of its close relative *R. speratus* (version 1) (19) as this gave better mapping results than attempting *de novo* genome assembly based on the short reads alone. From the resultant SAM files, the uniquely mapped reads were filtered using the Samtools tags (XS: i) (31). Male and female coverage was estimated using SoapCoverage (version 2.7.9; http://soap.genomics.org.cn/soapaligner.html). Coverage was summed up for all reads per scaffold to compare the coverage between males and females. Using R, the scaffolds were first filtered based on a minimum length of 7 kilobase-pairs (kb), followed by median normalisation to account for differences in the overall coverage of both sexes. This was followed by classifying scaffolds as either X-chromosomal (X) or autosomal (A), by calculating male-to-female (M:F) coverage and plotting coverage density histograms for visualisation. Scaffolds that had zero coverage in at least one sex were classified as unknown genomic location (‘NA’). The cutoff value on which scaffold classification was based was calculated based on the male-to-female ratio at the autosomal peak, which, due to differences in male and female coverage, was not always located directly at one. Distributions were thus shifted so that the autosomal peak was centered at one (Fig. 1 B). All scaffolds with a M:F coverage < 0.75 were defined as X-chromosomal while all scaffolds above this cut-off were defined as autosomal. In a more conservative approach, scaffolds with a M:F coverage < 0.6 were classified as X-chromosomal while the cutoff for the autosomal scaffold classification was maintained. In this more conservative approach, scaffolds that fell in between the two thresholds were classified as unknown (denoted as ‘NA’). The total size of the classified scaffolds was defined as the sum of all sex scaffold sizes and determined using a custom R script. As results did not qualitatively differ between the thresholds, we present here the former of the two while results for the more conservative approach can be found in the supplementary material.

### Gene expression analysis

To assess gene expression differences between males and females, RNA-sequencing (RNAseq) data of the separate sexes were used. Raw RNA-seq data of single male and female reproductives were downloaded from the NCBI Sequence Read Archive (SRA) for three cockroach species (*B. germanica, P. americana*, and *D. punctata*) and three termite species (*R. speratus, C. secundus*, and *M. natalensis*). For *B. germanica*, data stems from the three different projects BioProjects PRJNA780882, PRJNA38959, and PRJNA382128. Data of the remaining species is available under BioProject PRJNA779823, PRJDB5589, PRJNA707368, PRJNA382129, and PRJNA382034, respectively. For all species, except for *R. speratus*, whole body data sets were used. As whole body data was not available for *R. speratus*, two other composite tissue data sets were used: heads and thorax plus abdomen. For *B. germanica, D. punctata*, and the heads of *R. speratus*, three male and three female datasets were used. For *C. secundus*, four datasets were available per sex, while only two male and three female datasets were available for *M. natalensis, P. americana*, and *R. speratus* (thorax plus abdomen). Note that male RNAseq samples for *P. americana* came from two different strains and studies as no replication was avail-able in any single study. Coding sequences for the annotated genomes as well as protein sequences were downloaded for each species (see genome sources above).

### Read quality control and trimming

RNA-seq libraries composed of paired-end reads were quality checked using FASTQC (version 0.11.5; (32)) and subsequently trimmed using Trimmomatic (version 0.36; (33)), retaining only pairs of forward and reverse reads with a minimum of 36 bp after trimming. Furthermore, quality filtering of reads was performed using Trimmomatic sliding window which trims reads at the 5^*′*^ and 3^*′*^ ends, eliminating bases with a Phred score of < 15. For the short single-end reads of *R. speratus* and *D. punctata*, a sliding window function was used to remove fragments of four bases that had an average Phred score < 15, retaining only reads of 60bp or longer. Lastly, mapping was performed using NextGenMap (version 0.5.5; (34)) which aligned RNA-seq libraries to the coding-sequences of the respective genomes using default parameters.

### Sex-biased genes

To identify genes that are differentially expressed between the sexes of reproductives of each species, the Bioconductor package DESeq2 (version 1.10.1; (35)), as implemented in R, was used. The *R. speratus* analysis differs slightly from the other species as no whole body data was available. Instead, different tissue data sets, namely heads and thorax combined with abdomen were available and used here. As, especially for sex-biased genes in adult reproductives, we expect the major difference to be in the gonads, we present here the data for the combined thorax and abdomen samples. Nevertheless, all analyses have also been performed using heads.

Raw read counts obtained from the NextGenMap mappings (version 0.5.5; (34)) were used as the input and multiple testing correction was performed with the Benjamin-Hochberg correction as built into DESeq2. Genes with a significant adjusted P-value < 0.05 and a fold change *>* 2 between the sexes were considered sex-biased. All genes which did not meet this criterion were considered unbiased. Gene expression levels were measured as reads per kilobase of transcript per million mapped reads (RPKM) and unbiased genes with an average RPKM value < 0.5 were classified as being non-expressed.

### Gene Ontology (GO) enrichment analysis

The amino acid sequences download with the genomes for each species were used to annotate proteins via InterProScan (version 5.65-97.0; (36)). Of note, only the Pfam A database was used in InterProScan to transfer gene ontology (GO) terms as Pfam domains based on experimental validation were considered the most reliable. All other parameters were left as default. The Bioconductor package topGO (version 2.54.0; (37)) was used to assess over-represented GO terms among differentially expressed genes, taking the three individual GO categories (molecular function, cellular component and biological process) into account separately. Significant enrichment or deletion of terms was assessed in R with Fisher’s Exact Test using the Benjamini–Hochberg correction for multiple testing at a significance cutoff of an adjusted P-value < 0.05.

### Dosage Compensation Analysis

For dosage compensation analysis, gene expression levels were compared between the sexes and genomic locations (X-linked versus autosomal). As multiple isoforms were included in *C. secundus*, the mean expression of all isoforms was calculated per sex and used as gene expression value of downstream analysis. Quantile normalisation was performed for each individual using the normalize.quantiles function of the R Bioconductor pre-processCore package (version 1.66.0; (38)). Chromosomal assignment was performed using the results of the coverage-based analysis. Only genes with a mean RPKM *>* 1 in both sexes were kept for the downstream analysis. Replicates were averaged, followed by a second round of quantile normalisation. Using these RPKM values, gene expression was compared between (i) male and female autosomes, (ii) male and female X chromosomes, and (iii) male autosomes and the male X chromosome using Wilcoxon Rank Sum Test. Due to increased X-chromosomal gene expression in male individuals of *C. secundus* and *M. natalensis*, the analysis was repeated following exclusion of X-chromosomal genes with more than 2-fold expression in males compared to females.

### Sex-biased gene enrichment analysis

Quantile normalisation was performed and isoforms summarised as described above. All genes with a mean RPKM *>* 1 in at least one sex were included in the downstream analysis. Unbiased genes were defined as all genes with an adjusted P-value *>* 0.05 and sex bias was assigned for genes showing significantly different gene expression between sexes. The sex bias was assigned either solely based on significance (henceforth referred to as no cutoff, labelled as “none”) or each gene’s fold change between sexes was additionally considered for assignment. Two fold change cutoff values (fold change*>* 2 and *>* 4) were considered. To test for enrichment or depletion of sex-biased genes, the number of female-biased genes and male-biased genes on autosomes and the X chromosome was compared to the number of unbiased genes, respectively, using Fisher’s Exact Test (P-value < 0.05). The expected number of biased and unbiased genes on each chromosome type (autosomes and X chromosome) was extracted from each Fisher’s matrix and the ratio of observed-to-expected number of sex-biased genes was plotted for male- and female-biased genes on each chromosome type using the ggplot2 package (version 3.5.1; (39)).

### Across-species comparison

OrthoFinder (version 2.5.4; (40)) was used to filter for the longest isoform before determining orthogroups between all species (*B. germanica, P. americana, D. punctata, C. secundus, M. natalensis*, and *R. speratus*) by comparison of protein sequences. The OrthoFinder output was filtered for one-to-one (1:1) orthologues between all species and by each pairwise comparison. Furthermore, for pairwise comparison, 1:1 orthologues were also identified via Bidirectional Best Hit (BBH) BLAST (41) to gain more statistical power by individual comparisons. The overlap between sex-biased genes in the multispecies and pairwise comparison was visualised using the webtool Venny (version 2.0.2; (42)) and the ggvenn package (version 0.1.10) in ggplot2 (version 3.5.1; (39)) in R, respectively. Statistical significance of gene set overlaps was tested using the Super Exact (package SuperExactTest, version 1.1.0; (43)) and Fisher’s Exact Test for the multi-species and pairwise comparisons, respectively.

To assess if X-linked and autosomal genes were conserved between species, gene classifications determined from the coverage analysis were combined with the results of differential expression analysis and orthogroup predication data using dplyr (version 1.0.10; (44)) and ggplot2 (version 3.5.1; (39)) packages in R. The total number of genes that were X-linked or autosomal was determined in both species, followed by identification of genes which were autosomal in one species but X-linked in another and vice versa. Statistical testing was conducted to evaluate the association between these occurrences, using Fisher’s Exact Test. The number of genes overlapping between species as expected by chance was determined via Pearson’s Chi-squared Test. The analysis was repeated using the conservative sex scaffold classification.

## Results

### Differentiated X chromosome in one termite species

In diploid species with sex chromosomes, the heterogametic sex has only one of the shared sex chromosomes, namely one X chromosome in an XX/XY or XX/X0 system. Differentiation of sex chromosomes, including degeneration of the Y chromosome, reduces the number of reads for X-chromosomal scaffolds in the heterogametic sex. This allows for classification of sex scaffolds using coverage analysis. As genomes were mostly sequenced from females and Y chromosomes are often difficult to assemble especially with short reads, we have little to no sequence information for the Y chromosome. Our approach thus targeted the X chromosome to make inferences about its conservation across Blattodea. To identify the X chromosome in a unifying manner across Blattodea, three species of termites and three species of cockroaches were chosen for which genomes are currently available and which span the phylogeny of termites (Fig. 1 A). Wood roaches of the genus *Cryptocercus* would have furthermore been interesting to include but as currently no genome was available, they are not included in this analysis.

Autosomal and X-chromosomal origin was investigated using male-to-female coverage ratios of either published genomic data (for the cockroaches) or re-sequencing of males and females (for the termites). In each male-to-female coverage plot, the highest peak, centered at a coverage ratio of one, indicates equal coverage between males and females, as expected for autosomal regions (Fig. 1 B). The second highest peak, expected at coverage ratios less than one, corresponds to the X-chromosomal peak. These plots show that the male-over-female distributions are bimodal in the termite *M. natalensis* as well as in the German cockroach *B. germanica*, allowing for identification of X-linked scaffolds (Fig. 1 B). If the sexes are plotted separately, this is clearly due to the presence of a secondary X-chromosomal peak in males but not females (Fig. S1 A and B, supplementary material online). In contrast, the genomic coverage of both other termite species,

*R. flavipes* and *C. secundus*, shows only one peak (Fig. 1 B). In these species, both the male as well as the female coverage are unimodal with only an autosomal but no X-chromosomal peak clearly distinguishable (Fig. S1 C and D, supplementary material online). There is a small secondary peak in the male coverage of *C. secundus* but as there is also a stronger tail to the left of the autosomal distribution in the female (Fig. S1 D, supplementary material online), this does not lead to a clearly distinct peak in the male-to-female coverage plot (Fig. 1 B) and may simply be noise.

We also tried to identify X-linked scaffolds in two other species of cockroaches, namely in *P. americana* and *D. punctata*. However, as cockroaches have large genomes and the currently available data is still scarce, the coverage of these genomes proved too shallow for meaningful inferences. We nevertheless proceeded with these species as currently no better data is available but results are discussed in the light of these shortcomings. In *P. americana*, the only available dataset contained data of pooled males and females, while in *D. punctata*, only male data were available. In theory, a secondary peak indicating X-linked scaffolds could have been identified also in these datasets but with a mean coverage of 3.6 and 2.3 for *P. americana* and *D. punctata* respectively, possible identification of X-chromosomal scaffolds in both species was too limited (Fig. S1 E and F, supplementary material online). In *D. punctata*, the male coverage indicates that a secondary peak may be present but putative X-linked scaffolds (due to mixed sexes expected at 0.75 relative to the autosomal peak) were too close to zero to distinguish them from the noise caused by the many genomic positions with no or extremely low coverage (Fig. S1 F, supplementary material online).

### Larger X chromosome in cockroach than in termite

Depending on their male-to-female coverage, scaffolds were classified as autosomal (A), X-chromosomal (X), or unclassified (‘NA’), (Tab. S1, supplementary material online). In the German cockroach *B. germanica*, one third of all scaffolds were classified as X-chromosomal, corresponding to 245 mega base pairs (Mbp), approximately 12% of the whole genome size (Tab. S2, supplementary material online). In *P. americana*, approximately one fourth of the scaffolds from the genome assembly (namely 13 scaffolds) could be identified as X-linked. However, these only sum up to around 4 Mbp or 0.13% of this large genome. The situation is similar in *D. punctata*, where 8% of the scaffolds corresponding to 0.85% of the genome could be identified as X-linked in our analysis. In *M. natalensis*, the only termite species where we could identify a clearly differentiated X chromosome, over one fourth of all scaffolds were classified as X-chromosomal, corresponding to 82 Mbp or almost 7% of the whole genome. In *C. secundus* and *R. flavipes*, only 2.6% and 0.3% were classified as X-linked, respectively, analogous to the failure to detect a secondary peak indicating X-linkage (Fig. 1 B). Notably, in the more fragmented genomes of *M. natalensis* and *C. secundus*, many small scaffolds could not be classified but as these do not make up a large proportion of the genome, this is not expected to impact X chromosome size estimations strongly (Tab. S1 and Tab. S2, supplementary material online). Overall, as in the coverage analysis, only in the German cockroach *B. germanica* and the termite species *M. natalensis* could a large part of the genome be identified as X-linked. Here, the cockroach X chromosome is larger in absolute size but as cockroach genomes are larger than those of termites (2-3 giga base pairs (Gbp) versus 0.5-1 Gbp), this is expected. Nevertheless, the *B. germanica* X chromosome is also almost twice as large as the termite one relative to the genome size (12% versus 7% in *M. natalensis*).

For many of the species, the number of X-linked scaffolds relative to the total number of scaffolds in the genome assemblies and the absolute length of X-linked scaffolds is high indicating a higher degree of fragmentation in the genome assemblies for the X chromosome than for the autosomes.

### Sex chromosome turn-over within Blattodea

After classifying the scaffolds according to their X or autosomal linkage, we addressed the genes present on these scaffolds to test our three different hypotheses for the evolution of the XX/XY system in termites from the XX/X0 system in cockroaches. As expected based on the previous analysis, we found a large number of X-linked genes only in two species, namely *M. natalensis* and *B. germanica*, with over 1,000 genes in each of these and only 24-224 X-linked genes in the other species (Tab. 1).

**Table 1.** Total number of X-linked and autosomal genes in termite and cockroach species.

| Species | X-linked | Autosomal |
| --- | --- | --- |
| *B. germanica* | 1,710 | 27,869 |
| *P. americana* | 34 | 27,013 |
| *D. punctata* | 180 | 28,234 |
| *C. secundus* | 224 | 17,998 |
| *R. speratus* | 35 | 15,556 |
| *M. natalensis* | 1,267 | 15,042 |

We thus concentrated in our comparison on the termite *M. natalensis* and the German cockroach *B. germanica* which are the two species where coverage analysis revealed a clear secondary peak for X-linked scaffolds (Fig. 1 B), indicating that the X chromosomes of both species is rather large and differentiated. We identified orthologues between the species and found only four genes to be X-chromosomal in both species while almost all of X-chromosomal genes in each species (287 in *M. natalensis* and 367 in *B. germanica*) are autosomal in the other species (Tab. 2). This overlap is smaller than expected (FET, *P* = 0.003). Interestingly, of the four conserved X-linked genes (Tab. S3, supplementary material online), two are located on the same scaffold in *B. germanica* but on different scaffolds in *M. natalensis*. Both of these genes were classified as X-linked also with the more conservative parameters, further increasing reliability (overlap also smaller than expected by chance, FET, *P* = 0.026). This comparative analysis shows that the majority of genes which are X-linked in one species is autosomal in the other and vice versa, indicating sex chromosome turn-over within Blattodea (Tab. 2).

**Table 2.** Numbers of X-linked and autosomal genes shared between the termite *M. natalensis* and the cockroach *B. germanica*.

|  |  | *B. germanica* |  |
| --- | --- | --- | --- |
|  |  | X-linked | Autosomal |
| *M. natalensis* | X-linked | 4 | 287 |
|  | Autosomal | 367 | 7179 |

Each of these genes with concurrent X linkage in a termite and a cockroach were blasted to identify possible sequence similarities with other species. Results show that one gene pair (*Mnat_17008* and *Bger_01236*) has no homology to known genes on NCBI, while the *B. germanica* gene on the same scaffold (*Bger_01233*) is homologous to genes from the *yellow* gene family and shares sequence similarity with another termite species, *Zootermopsis nevadensis*. The other two genes show homology to other insects and are similar to genes annotated as *snakeskin* and *GrpE protein-like* in other social insects and *brachyurin*, a serine-endopeptidase. For two of these X-linked gene pairs, GO terms could be annotated and they are mostly associated with functions related to protein metabolism such as folding, dimerisation, binding, trafficking, and proteolysis (Tab. S4, supplementary material online).

To test whether there could be at least a small conserved X-linked region that is present across termite species, we assessed the overlap between genes identified as X-linked in *M. natalensis*, where we can identify the secondary peak in the coverage distribution, and the other two species *C. secundus* and *R. flavipes*, where we cannot. Despite the low numbers of X-linked genes, statistical analysis using Fisher’s Exact Test and Pearson’s Chi-squared Test revealed a significant positive association in the overlap of X-linked genes between *R. flavipes* and *C. secundus* with *M. natalensis* (*P* = 0.008 and *P* = 0.011, respectively; Tab. S5 and S6, supplementary material online). These results indicate that the number of X-linked genes overlapping between these species is higher than expected by chance, suggesting that at least a small part of the X chromosome may be conserved within the group of termites. As with four and three genes, however, this is not (at least in terms of protein-coding genes) larger than what we find conserved between the termite and the cockroach.

### Fast turn-over of sex-biased genes across Blattodea

To further assess sex differences and characterise the X chromosome in more detail, gene expression between the sexes was analysed and compared across species. In order to compare sex-biased gene expression in the solitary cockroaches and the social termites, only male and female termites of the reproductive caste, where sexual dimorphism is most pronounced, were used in this study. In the termite *R. flavipes*, no current genome is available and for the genomic analysis, reads were thus mapped to the closely-related sister species *R. speratus*. For this species, also RNAseq data was available and hence the gene expression study was performed with this species rather than *R. flavipes*.

As pointed out before, cockroaches possess larger genomes overall but also a higher number of protein-coding genes (approx. 27,000-30,000 compared to approx. 15,000-18,000 in termites). Overall, the cockroaches *B. germanica* and *D. punctata*, with the exception of *P. americana*, displayed a greater number of sex-biased genes than termites (absolute and relative to proteome size), with sex-biased genes accounting for one forth of all genes (25.12% and 25.44%, respectively, Fig. 2 A and Tab. S7, supplementary material online). In *P. americana*, only 1.77% of all genes show sex-biased expression. However, this is most probably due to the fact that RNAseq data from multiple studies (originating from different strains in the male samples) had to be pooled because no biological replicates would have been available otherwise. These data showed large variability in gene expression profiles and thus these few sex-biased genes are probably the most extreme ones that are also conserved across different strains. This is further reflected by the fact that almost all of these 480 genes (namely 476) show extremely strong sex bias with a more than four-fold expression difference between the sexes as well as a very high number of genes that were classified as “non-expressed” in our analysis as fold changes and P-values could not be calculated in DESeq2 due to strong dispersion of values (Fig. 2 A). Hence, for a more comprehensive study in the cockroach *P. americana*, more and better data is necessary.

**Fig. 2.**
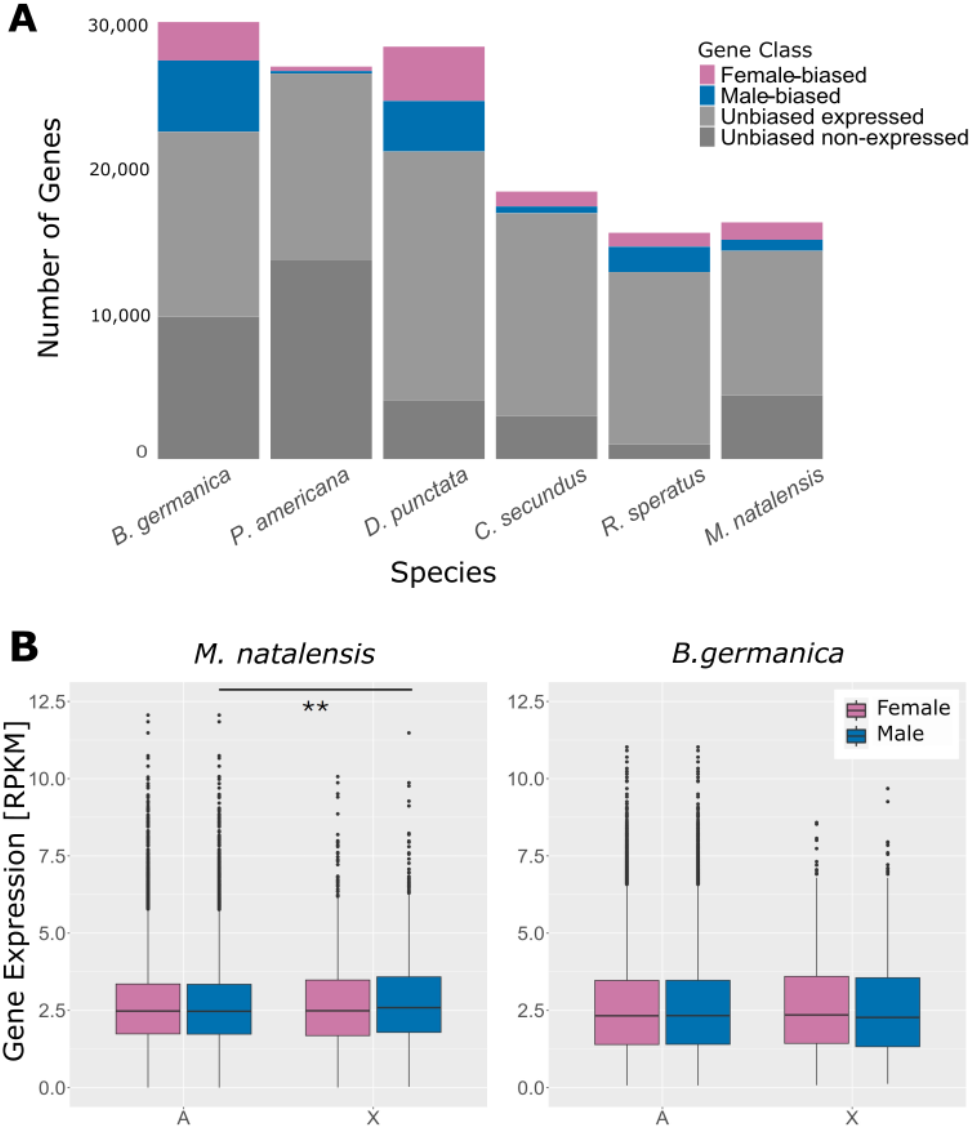
Gene expression differences between the sexes in Blattodea. (A) Numbers of genes showing female-biased (pink), male-biased (blue), and unbiased expression (grey) were determined across termite and cockroach species. Unbiased genes were further divided into clearly unbiased genes which are expressed (light grey) as well as unbiased genes which are non-expressed or too variable between replicates to calculate the relevant statistics (dark grey). (B) Dosage compensation in whole body gene expression data balances X chromosomal expression in male *M. natalensis* and *B. germanica*. Normalised X-chromosomal (X) and autosomal (A) gene expression values was plotted as RPKM values for females (pink) and males (blue) of the termite *M. natalensis* and the cockroach *B. germanica*. Statistical analysis was performed using Wilcoxon Rank Sum Test; * *P* < 0.05; ** *P* < 0.01, *** *P* < 0.001.

In the other two cockroaches, 8.80% and 13.17% are female-biased, while 16.32% and 12.27% are male-biased, in *B. germanica* and *D. punctata* respectively. Sex-biased genes in *R. speratus, M. natalensis*, and *C. secundus* correspond to only 17.37% (6.15% female-biased and 11.22% male-biased), 12.06% (7.37% female-biased, 4.69% male-biased), and 8.11% (5.57% female-biased, 2.53% male-biased) of genes, respectively (Fig. 2 A and Tab. S7, supplementary material online). Furthermore, across Blattodea, there is no clear pattern, like in some other species, that male-biased genes are more numerous than female-biased genes. However, it is important to notice that for most of these species (except for *R. speratus*) only whole body data is currently available and as tissue sizes differ between the species, allometric scaling may affect these numbers. Hence, more detailed studies about the evolution of sex-biased genes would require profiling individual tissues. In summary, the total number but also the percentage of genes with differential expression between the sexes is higher in cockroaches than in termites (Fig. 2 A and Tab. S7, supplementary material online).

To identify prominent biological function of sex-biased genes, Gene Ontology (GO) analysis was performed, revealing between 14 and 43 GO terms enriched in the sex-biased genes for each species. Enriched GO terms relate to a variety of different function and can be found in Tab. S8, supplementary material online. Interestingly, no sex-biased genes and also no functional GO terms were conserved amongst all species or within the termites. However, a total of seven GO terms were conserved amongst the cockroaches. These conserved GO terms are all related to serine-type endopeptidase activities (Tab. S9, supplementary material online). Interestingly, all GO terms were female-biased in *B. germanica* and *P. americana* but male-biased in *D. punctata*. In summary, these results indicate no conserved function of sex-biased genes within Blattodea and overall fast evolution of sex-biased gene expression.

### Enrichment of sex-biased genes on the X chromosome of *M. natalensis* and *B. germanica*

Sex chromosomes are known to accumulate sex-biased genes which, in the case of male heterogamety, often leads to feminisation of the X chromosome (45, 46) in line with theoretical predictions (47). Furthermore, enrichment of strongly sex-biased genes on the sex chromosomes can obscure patterns of dosage compensation (48). To investigate potential enrichment of sex-biased genes on the X chromosome, the number of female- and male-biased genes on autosomes and X chromosomes were compared to the number of unbiased genes, respectively, using Fisher’s Exact Test. Additionally to using all genes identified as differentially expressed by adjusted P-value alone, two different cutoffs (2- and 4-fold difference between males and females) were used to investigate whether the strength of sex bias affects enrichment or depletion of said genes. Below, the cutoff “none” refers to all sex-biased genes being included into the analysis, independently of their fold change between sexes. Notably, no sex-biased genes were identified in the head of *R. speratus* which was therefore excluded from further analyses. In this section, all data referring to *R. speratus* was obtained from the thorax and abdomen samples.

As expected, results are clearest in those two species for which we could identify a significant amount of X-linked genes, namely *M. natalensis* and *B. germanica*. Interestingly, almost all genes show rather pronounced sex bias with a fold change difference between males and females of at least two (Tab. S10, S11, and S12, supplementary material online). Smaller differences between the sexes can be attributable to a lack of dosage compensation as often seen in gonads (48–51). In the termite *M. natalensis*, we find male-biased genes to be significantly enriched on the X chromosome, independent of their strength of sex bias (FET, *P* < 0.001 in all comparisons; Fig. S4, supplementary material online), while only strongly female-biased genes are enriched on the X chromosome (*P*_4_ = 0.0046, supplementary material online; n_*X−linkedGenes*_ = 880; Tab. S10, Fig. S4 C, supplementary material online). The other two termite species, *R. speratus* and *C. secundus* generally have very few genes on few scaffolds that are presumably X-linked (Tab. 1) and of those, only between 7 and 9 are sex-biased (Tab. S10, supplementary material online) making an inference difficult. *C. secundus* shows a slight depletion of female-biased genes but not for highly female-biased genes (*P*_*none*_ = 0.03, *P*_2_ = 0.03, *P*_4_ = 0.73; Fig. S4 and S5, supplementary material online). Quantification of sex-biased genes in the German cockroach *B. germanica* reveals a significant enrichment of female-biased genes on the X chromosome independent of their strength of sex bias (*P*_*none*_ = 0.02, *P*_2_ = 0.003, *P*_4_ = 0.0005, supplementary material online), suggesting feminisation of the X chromosome. The number of sex-biased genes on the X chromosome was neither significantly enriched nor depleted in the other two cockroaches (*P >* 0.05 for all comparisons) but of course numbers of X-linked genes and thus statistical power was much lower for these species.

### Full dosage compensation of the X chromosomes of Blattodea

In order to assess whether dosage compensation is present in Blattodea, we compared the gene expression levels between the X-linked and the autosomal genes in males and females of the different species. We performed the analysis for all species involved (see methods and Fig. 1 A) but chose to show here the two species which we find to have differentiated X chromosomes. Overall, the analysis revealed no significant differences between male and female X chromosomes in either species, suggesting balanced gene expression between sexes (Fig. 2 B and Fig. S2 A, supplementary material online). This pattern consistently holds up using the more conservative sex chromosome scaffold classification (Fig. S3 A, supplementary material online).

The comparison of gene expression between the male autosomes and the male X chromosome revealed no significant differences in the termite *R. speratus* and the cockroaches *D. punctata, B. germanica*, and *P. americana* (Fig. S2 A, supplementary material online).

Interestingly, the expression of X-linked genes was significantly higher than that of autosomal genes in males of *M. natalensis* and *C. secundus* (*P* = 0.003 and *P* = 0.002, respectively, Fig. 2 B and Fig. S2 A, supplementary material online). In *M. natalensis*, significantly higher expression of X-chromosomal compared to autosomal genes was confirmed using the conservative scaffold classification (Fig. S3 A, supplementary material online). This pattern may be a result of enrichment of male-biased genes on the X chromosome as has been reported in other species (48). To test this potential explanation, X-chromosomal genes with more than 2-fold expression in males compared to females were excluded before repeating the analysis in *M. natalensis*. Using this approach, the patterns largely hold but the difference in X and autosomal expression in males disappears (*P* = 0.971, Fig. S2 B, supplementary material online). This suggests that dosage compensation is indeed complete and only the slightly higher expression of the X chromosome in males than the autosomes can be attributed to an enrichment of male-biased genes on the X chromosome of *M. natalensis*. In *C. secundus*, no male-biased genes with more than two-fold expression difference were found on the X chromosome (Tab. S11, supplementary material online). It is therefore possible that an overshoot in dosage compensation mechanisms might be responsible for increased male X-chromosomal gene expression.

In summary, the combination of equal gene expression of the X chromosomes between sexes and the expression of X-chromosomal genes at the same or even at a higher level than autosomal genes in males suggests full dosage compensation to have evolved in cockroaches as well as termites.

## Discussion

We present here the first comparative study using genomic and transcriptomic data of the sex chromosomes and their evolution across Blattodea to which termites, together with cockroaches, belong. While sex determination is based on a male-heterogametic system throughout this order, no Y chromosome is present in the cockroaches, leading to a XX/X0 system, while termites possess a XX/XY system (5–7). Phylogenetically, termites have evolved from within the paraphyletic cockroaches (8, 9), leading to the assumption that the XX/X0 system of cockroaches is the ancestral state in Blattodea. This is the most parsimonious explanation as otherwise the Y chromosome would have to be lost in all branches of cockroaches independently. Assuming an ancestral XX/X0 systems raises the question where the Y chromosome came from. Three scenarios can explain the Y chromosome gain: (i) turn-over of sex chromosomes, (ii) chromosomal fusion of the ancestral X chromosome to an auto-some, resulting in the fused chromosome as a neo-X chromosome and the homologous unfused chromosome as a neo-Y chromosome, and (iii) recruitment of an extra-genomic element (27). Each mechanism would result in a different hypothetical outcome regarding differentiation, homology, and sex chromosome size, as discussed above. Based on these hypotheses, we here provide evidence for scenario (i) suggesting complete turn-over with a novel pair of differentiated sex chromosomes in at least one termite species.

### The termite *M. natalensis* possesses large, differentiated sex chromosomes

In this study, we studied size, gene content, and gene expression of the X chromosomes of multiple species of cockroaches and termites in order to deduce how the sex chromosomes evolved in Blattodea. We used a coverage-based approach to test whether differentiated sex chromosomes are present. Within Blattodea, dual-peaked distributions, resulting from the presence of only one X chromosome in males, can only be detected in the termite *M. natalensis* and the cockroach *B. germanica*. These X-linked scaffolds represent a large part of the genome with the X chromosome being larger in the cockroach (12% versus 7% in the termite), indicating large and differentiated X chromosomes in both of these species. This result is in line with what has been found in *B. germanica* previously (17). In contrast, no clear secondary peak was detected in the other two termites *C. secundus* and *R. flavipes* (mapped to the *R. speratus* genome due to a lack of data on this species), indicating that their X chromosome may not be a as large and differentiated as in the first two species. It is possible that these sex chromosomes are evolutionary younger with less degeneration of the Y chromosome. This would mean that reads from the Y chromosome can still map to X-linked scaffolds such that a difference in coverage would not be easily detectable in our analysis. According to Jankasek et al. (7), these two species have not been investigated in depth yet but due to presence of XX/XY sex determination in closely-related species of the same genus, an XX/XY system can also be assumed for them (52–54). Thus, it is also possible that these species have alternative systems of sex determination not including differentiated sex chromosomes such as polygenic sex determination. However, in *R. speratus*, multivariate chains have been described that usually include the sex chromosomes (54) and the same is true for *R. flavipes* (McMahon, personal communications) suggesting that these species do indeed possess sex chromosomes.

Alternatively, more technical issues may hinder identification of a differentiated X chromosome in those two species. Our analyses indicate that overall the X chromosomal scaffolds are more numerous and shorter than for the autosomes. This is not surprising as sex chromosomes are often hard to assemble due to the extensive presence of transposable elements and repeats (55–57) and only now do we start to have full-length sex chromosome assemblies for a handful of model organisms (58–61). Thus, gaps or errors in the current genome assemblies may hinder a clear identification of X-linked scaffolds. In addition, we only included scaffolds of a certain size (7 kbp here, but also tested with lower thresholds such as 2 or 4 kbp) to reduce noise and, especially if the X chromosome is very fragmented in the assemblies, some of the signal may have been lost.

High quality genomes with full chromosome-level assemblies for Blattodea will allow further investigation of the sex chromosomes across many species including the detection of different strata which with the current genome assemblies is not possible. Nevertheless, our results clearly indicate variability in terms of size and level of differentiation of the X chromosome within the termites which makes this group very interesting for further studies into sex chromosome and genome structure evolution.

### Lacking homology between the termite and cockroach X chromosomes suggests complete turn-over

Comparative analysis of the gene content of the termite *M. natalensis* with the German cockroach *B. germanica* demonstrates little homology between their respective X chromosomes. In fact, almost all X-linked genes of *B. germanica* are autosomal in the termite *M. natalensis* and vice versa, suggesting complete turn-over of sex chromosomes. This supports a scenario where the X chromosome of cockroaches has reverted back to an autosome in termites and a new pair of sex chromosomes has been recruited. Fast and recurrent turn-overs of sex chromosomes, even in a dense phylogenetic context, have been described before, for example in *Drosophila* (49, 62, 63). It is possible that the special biology of termites with its recurrent occurrence of bivalent chains that include the sex chromosomes may facilitate fast karyotypic evolution (64, 65) rendering Blattodea an especially interesting group to study sex chromosome evolution.

Only four genes are found to be X-linked in both species, two of them on the same scaffold in the cockroach genome. It is possible that these genes form part of an ancestral sex determining region that may still be conserved across Blattodea. The presence of such a region, at least within termites, is supported by the fact that the overlap of X-linked genes in the termite *M. natalensis* with the other two termites is larger than expected by chance. As discussed above, we may have missed the differentiated sex chromosomes of these two species in our analysis due to current data limitations. Alternatively, more sex chromosome turn-over events may have taken place within termites potentially as the result of a translocation of the sex-determining region. Such translocations of existing master sex-determining genes are, together with the evolution of a novel master sex determining gene, one of the best documented causes for sex chromosome turn-over (reviewed in (26)). The evolution of novel master sex determining genes that compete with existing ones has also been found in species such as the house fly, a system that is believed to be currently in transition (66). Relocation of a sex determining region, on the other hand, has been found in several other taxa using synteny approaches and whole genome alignments (67–69). More complete and contiguous genomes will make these kinds of analyses possible and thus help elucidate the way sex chromosome turn-overs happened in termites.

This study clearly shows that sex chromosome turn-over (scenario i) has occurred between the XX/X0 system of cockroaches and the XX/XY system of termites. This is further supported by the fact that the X chromosome is larger in the cockroach than in the termite. In case of a fusion of the cockroach X to an autosome, we would have expected cockroach X-linked genes to be largely X-linked in the termites as well and for the X chromosome to have increased in size. Of course other events such as translocations, in- or decrease of repeat content, etc. also play important roles over these long evolutionary time scales. However, the fact that the cockroach X chromosome has been found to be an ancestral X across insects and homology can be detected throughout the whole class (17, 18) shows that these process may be of minor importance for chromosome-wide patterns.

### X chromosomes display typical hallmarks of a sex chromosome

Evolution of sex chromosomes is often accompanied by accumulation of sex-biased and sex-specific genes on the sex chromosomes (reviewed in (70, 71)). In male-heterogametic systems, this often leads to feminisation of the X chromosome (45, 46). Termites have an overall surprisingly low number of sex-biased genes, given that we included only reproductives and whole body data that include the gonads, the main source of gene expression differences between the sexes. Most other species such as *Drosophila* show a greater proportion of genes with sex-biased expression (50). The proportions of sex-biased genes in the cockroaches are more similar to what we find in other insects. As we used publicly available gene expression data here, we cannot determine whether individuals used in these different species were actually reproductively active. Nevertheless, we show that sex-biased genes are indeed enriched on the X chromosomes of the termite *M. natalensis* that displays differentiated sex chromosomes. This is also true for the cockroach *B. germanica*, where the X chromosome shows pronounced feminisation of the X chromosome. This is consistent with observations in other species with evolutionarily old and highly differentiated sex chromosomes. In the termite *M. natalensis*, however, both types male- and female-biased genes are enriched on the X chromosome. As our data includes gonads, it is possible that a strong signal in the males comes from this tissue, which often harbours many very strongly-biased genes. More fine-scale analysis of individual tissue, especially meiotic and somatic tissues separated, will be necessary to disentangle this, stressing the need for more and more comparable data on multiple termite species.

Differentiation of sex chromosomes, characterised by degeneration of the sex-limited chromosome, leads to gene dosage imbalance in the heterogametic sex. Thus, differentiated sex chromosomes often show by mechanisms balancing transcription of sex chromosomes that restore X-linked gene expression ((72) and reviewed in (73, 74)). In XX/XY or XX/X0 species with full dosage compensation, the male X chromosome expression matches the expression of diploid autosomes, leading to an X-to-autosome (X:A) expression ratio of one and equal expression of the X chromosome in the two sexes. Dosage balance refers to equalised expression only between the sexes but without an X-to-autosome ratio of one (51). Our study shows that the X chromosomes of termites and cockroaches seem to be fully dosage compensated with mostly equal expression levels of the X chromosome and autosomes in males and equal expression of X-linked genes between males and females. In *B. germanica*, first analyses had also indicated full X chromosome dosage compensation, but final conclusions were hindered by the fact that only mixed sex data and different tissue for the different sexes had been available (17). In the termites, we observe even a slightly higher expression of X-linked genes in the males than for autosomal genes. As this increase is insignificant when excluding X-chromosomal male-biased genes, it may be caused by enrichment of male-biased genes on the X chromosome of *M. natalensis*. In *C. secundus*, on the contrary, significance is not reduced when excluding said genes and increased X-chromosomal gene expression may thus be due to overshooting dosage compensation as seen in *Drosophila* where this also varies between tissues (50), further highlighting the need to obtain more samples from different tissue and developmental stages to study gene regulation in termites in closer detail. Overall, we see presence of dosage compensation and accumulation of sex-biased genes, not only on the X chromosome of *B. germanica*, which has been confirmed to be evolutionarily old (17, 18), but also on the X chromosome of the termite *M. natalensis* that we newly identified and described in this study.

## Conclusions

We here provide evidence for sex chromosome turn-over, accompanied by pronounced differentiation, between cockroaches and at least one termite species, namely *M. natalensis*. The overall pattern supports complete sex chromosome turn-over, where the XX/XY system of termites evolved from an XX/X0 system still present in cockroaches and woodroaches. The ancestral X chromosome has reverted back to an autosome in termites and a new pair of autosomes has formed the X and Y chromosomes. As expected under this scenario, hardly any conservation of X-linked genes is observed in the comparison of termites to cockroaches. The X chromosome in this termite species displays typical hallmarks of evolutionarily old sex chromosomes. As such, the X chromosome is enriched for sex-biased genes and complete dosage compensation is observed, as is also the case in the German cockroach *B. germanica*. While conservation of some genes on sex-linked scaffolds within termites suggests that at least a part of the novel X chromosome, potentially the sex determining region, is shared between these species, this needs further confirmation using improved and more contiguous genome assemblies.

## Author Contributions

AKH conceived the study and acquired the funding. AKH and BV designed the study and performed the sequencing. AKH, AD, RF, RM, NM, and SL analysed the data. AKH, AD, RF, and RM wrote the manuscript. All authors have read and approved the manuscript.

## ACKNOWLEDGEMENTS

We thank all lab members and collaborators for feedback on the project. Specifically, Dino McMahon provided *R. flavipes* males and females, Judith Korb provided *C. secundus* males and females, gave feedback on the project and discussed questions on termite reproduction, Mireille Vasseur-Cognet provided *M. natalensis* males and females, Ariana Macon performed the lab work for sequencing and the Vicoso group gave critical feedback on the project. We furthermore thank the HPC group at IST Austria and Christian Meesters at JGU Mainz for their technical support.

## Bibliography

1. C. Noirot. Recherches sur le polymorphisme des termites supérieurs (termitidae). Ann. Sci. Nat. Zool., 17:399–595, 1955.

2. Yves Roisin. Intragroup conflicts and the evolution of sterile castes in termites. The American Naturalist, 143:751–765, May 1994. ISSN 0003-0147. Submitted February 18, 1993; Revised June 30, 1993; Accepted June 26, 1993.

3. Judith Korb and Klaus Hartfelder. Life history and development -a framework for understanding developmental plasticity in lower termites, 8 2008. ISSN 14647931.

4. Mark C. Harrison, Evelien Jongepier, Hugh M. Robertson, Nicolas Arning, Tristan Bitard-Feildel, Hsu Chao, Christopher P. Childers, Huyen Dinh, Harshavardhan Doddapaneni, Shannon Dugan, Johannes Gowin, Carolin Greiner, Yi Han, Haofu Hu, Daniel S.T. Hughes, Ann Kathrin Huylmans, Carsten Kemena, Lukas P.M. Kremer, Sandra L. Lee, Alberto Lopez-Ezquerra, Ludovic Mallet, Jose M. Monroy-Kuhn, Annabell Moser, Shwetha C. Murali, Donna M. Muzny, Saria Otani, Maria Dolors Piulachs, Monica Poelchau, Jiaxin Qu, Florentine Schaub, Ayako Wada-Katsumata, Kim C. Worley, Qiaolin Xie, Guillem Ylla, Michael Poulsen, Richard A. Gibbs, Coby Schal, Stephen Richards, Xavier Belles, Judith Korb, and Erich Bornberg-Bauer. Hemimetabolous genomes reveal molecular basis of termite eusociality. Nature Ecology and Evolution, 2:557–566, 3 2018. ISSN 2397334X. doi: 10.1038/s41559-017-0459-1.

5. P.P. Vincke and J.P. Tilquin. A sex-linked ring quadrivalent in termitidae (isoptera). Chromosoma, 67:151–156, June 1978. doi: 10.1007/BF00293172. Received 05 November 1977; Accepted 05 November 1977.

6. Silvia Bergamaschi, Tracy Z. Dawes-Gromadzki, Valerio Scali, Mario Marini, and Barbara Mantovani. Karyology, mitochondrial dna and the phylogeny of australian termites. Chromosome Research, 15:735–753, 10 2007. ISSN 09673849. doi: 10.1007/s10577-007-1158-6.

7. Marek Jankásek, Zuzana Kotyková Varadínová, and František Št’áhlavský. Blattodea karyotype database. European Journal of Entomology, 118, 2021. ISSN 18028829. doi: 10.14411/EJE.2021.020.

8. Daegan Inward, George Beccaloni, and Paul Eggleton. Death of an order: a comprehensive molecular phylogenetic study confirms that termites are eusocial cockroaches. Biology letters, 3(3):331–335, 2007.

9. Nathan Lo, Gaku Tokuda, Hirofumi Watanabe, Harley Rose, Michael Slaytor, Kiyoto Maekawa, Claudio Bandi, and Hiroaki Noda. Evidence from multiple gene sequences indicates that termites evolved from wood-feeding cockroaches. Current Biology, 10:801–804, June 2000. doi: 10.1016/S0960-9822(00)00571-X. Received: 15 February 2000; Revised: 27 March 2000; Accepted: 25 May 2000; Published: 16 June 2000.

10. M.J.D. White. Cytogenetics of orthopteroid insects. In M. Demerec, editor, Advances in Genetics, volume 4 of Advances in Genetics, pages 267–330. Academic Press, 1951. doi: 10.1016/S0065-2660(08)60238-2.

11. Donald G. Cochran and Mary H. Ross. Preliminary studies of the chromosomes of twelve cockroach species (blattaria: Blattidae, blattellidae, blaberidae). Annals of the Entomological Society of America, 60(6):1265–1272, November 1967. doi: 10.1093/aesa/60.6.1265.

12. Srinivas Kambhampati, Peter Luykxt, and Christine A Nalepa. Evidence for sibling species in cryptocercus punctulatus, the wood roach, from variation in mitochondrial dna and karyotype, 1996.

13. Nathan Lo, Peter Luykx, Rossana Santoni, Tiziana Beninati, Claudio Bandi, Maurizio Casiraghi, Lu Wen-Hua, Evgueni V. Zakharov, and Christine A. Nalepa. Molecular phylogeny of cryptocercus wood-roaches based on mitochondrial coii and 16s sequences, and chromosome numbers in palearctic representatives. Zoological Science, 23:393–398, 4 2006. ISSN 02890003. doi: 10.2108/zsj.23.393.

14. Christine A. Nalepa, Keisuke Shimada, Kiyoto Maekawa, and Peter Luykx. Distribution of karyotypes of the cryptocercus punctulatus species complex (blattodea: Cryptocercidae) in great smoky mountains national park. Journal of Insect Science, 17, 5 2017. ISSN 15362442. doi: 10.1093/jisesa/iex045.

15. Thomas Bourguignon, Nathan Lo, Stephen L Cameron, Jan Šobotník, Yoshinobu Hayashi, Shuji Shigenobu, Dai Watanabe, Yves Roisin, Toru Miura, and Theodore A Evans. The evolutionary history of termites as inferred from 66 mitochondrial genomes. Molecular biology and evolution, 32(2):406–421, 2014.

16. Ales Bucek, Jan Šobotník, Shulin He, Mang Shi, Dino P McMahon, Edward C Holmes, Yves Roisin, Nathan Lo, and Thomas Bourguignon. Evolution of termite symbiosis informed by transcriptome-based phylogenies. Current Biology, 29(21):3728–3734, 2019.

17. Richard P. Meisel, Pablo J. Delclos, and Judith R. Wexler. The x chromosome of the german cockroach, blattella germanica, is homologous to a fly x chromosome despite 400 million years divergence. BMC Biology, 17, 12 2019. ISSN 17417007. doi: 10.1186/s12915-019-0721-x.

18. Clementine Lasne, Marwan Elkrewi, Melissa A. Toups, Lorena Layana, Ariana Macon, an Beatriz Vicoso. The scorpionfly (panorpa cognata) genome highlights conserved and derived features of the peculiar dipteran x chromosome. Molecular Biology and Evolution, 40, 12 2023. ISSN 15371719. doi: 10.1093/molbev/msad245.

19. Shuji Shigenobu, Yoshinobu Hayashi, Dai Watanabe, Gaku Tokuda, Masaru Y Hojo, Kouhei Toga, Ryota Saiki, Hajime Yaguchi, Yudai Masuoka, Ryutaro Suzuki, Shogo Suzuki, Moe Kimura, Masatoshi Matsunami, Yasuhiro Sugime, Kohei Oguchi, Teruyuki Niimi, Hiroki Gotoh, Masaru K Hojo, Satoshi Miyazaki, Atsushi Toyoda, Toru Miura, and Kiyoto Maekawa. Genomic and transcriptomic analyses of the subterranean termite reticulitermes speratus: Gene duplication facilitates social evolution. PNAS, 119:e2110361119, 2022. doi: 10.1073/pnas.2110361119/-/DCSupplemental.

20. Jacopo Martelossi, Giobbe Forni, Mariangela Iannello, Castrense Savojardo, Pier Luigi Martelli, Rita Casadio, Barbara Mantovani, Andrea Luchetti, and Omar Rota-Stabelli. Wood feeding and social living: Draft genome of the subterranean termite reticulitermes lucifugus (blattodea; termitoidae). Insect Molecular Biology, 32:118–131, 4 2023. ISSN 13652583. doi: 10.1111/imb.12818.

21. Michael Poulsen, Haofu Hu, Cai Li, Zhensheng Chen, Luohao Xu, Saria Otani, Sanne Nygaard, Tania Nobre, Sylvia Klaubauf, Philipp M Schindler, Frank Hauser, Hailin Pan, Zhikai Yang, Anton S M Sonnenberg, Z Wilhelm De Beer, Yong Zhang, Michael J Wingfield, J P Grimmelikhuijzen, Ronald P De Vries, Judith Korb, Duur K Aanen, Jun Wang, Jacobus J Boomsma, and Guojie Zhang. Complementary symbiont contributions to plant decomposition in a fungus-farming termite. PNAS, 11:14500–14505, 2014. doi: 10.5524/100055.

22. Nicolas Terrapon, Cai Li, Hugh M. Robertson, Lu Ji, Xuehong Meng, Warren Booth, Zhensheng Chen, Christopher P. Childers, Karl M. Glastad, Kaustubh Gokhale, Johannes Gowin, Wulfila Gronenberg, Russell A. Hermansen, Haofu Hu, Brendan G. Hunt, Ann Kathrin Huylmans, Sayed M.S. Khalil, Robert D. Mitchell, Monica C. Munoz-Torres, Julie A. Mustard, Hailin Pan, Justin T. Reese, Michael E. Scharf, Fengming Sun, Heiko Vogel, Jin Xiao, Wei Yang, Zhikai Yang, Zuoquan Yang, Jiajian Zhou, Jiwei Zhu, Colin S. Brent, Christine G. Elsik, Michael A.D. Goodisman, David A. Liberles, R. Michael Roe, Edward L. Vargo, Andreas Vilcinskas, Jun Wang, Erich Bornberg-Bauer, Judith Korb, Guojie Zhang, and Jürgen Liebig. Molecular traces of alternative social organization in a termite genome. Nature Communications, 5, 5 2014. ISSN 20411723. doi: 10.1038/ncomms4636.

23. Shuji Itakura, Yuya Yoshikawa, Yasuhiro Togami, and Kiwamu Umezawa. Draft genome sequence of the termite, coptotermes formosanus: Genetic insights into the pyruvate dehydrogenase complex of the termite. Journal of Asia-Pacific Entomology, 23:666–674, 8 2020. ISSN 12268615. doi: 10.1016/j.aspen.2020.05.004.

24. Adam S Wilkins. bies.950170113. BioEssays, 17:71–77, 1995.

25. Andrew Pomiankowski, Rolf Nöthiger, Adam Wilkins, and Stephenson Way. The evolution of the drosophila sex-determination pathway studies with other species, 2004.

26. Benjamin LS Furman, David CH Metzger, Iulia Darolti, Alison E Wright, Benjamin A Sandkam, Pedro Almeida, Jacelyn J Shu, and Judith E Mank. Sex chromosome evolution: so many exceptions to the rules. Genome biology and evolution, 12(6):750–763, 2020.

27. Christelle Fraïsse, Marion A.L. Picard, and Beatriz Vicoso. The deep conservation of the lepidoptera z chromosome suggests a non-canonical origin of the w. Nature Communications, 8, 12 2017. ISSN 20411723. doi: 10.1038/s41467-017-01663-5.

28. Ben Langmead and Steven L. Salzberg. Fast gapped-read alignment with bowtie 2. Nature Methods, 9:357–359, 4 2012. ISSN 15487091. doi: 10.1038/nmeth.1923.

29. Lingyi Wang, Qing Xiong, Nawannaporn Saelim, Lin Wang, Wenyan Nong, Angel Tsz-Yau Wan, Mai Shi, Xiaoyu Liu, Qin Cao, Jerome Ho Lam Hui, Nitat Sookrung, Ting-Fan Leung, Anchalee Tungtrongchitr, and Stephen Kwok Wing Tsui. Genome assembly and annotation of periplaneta americana reveal a comprehensive cockroach allergen profile. Allergy, 78(4):1088–1103, Apr 2023. doi: 10.1111/all.15531. Epub 2022 Oct 5.

30. Benjamin Fouks, Mark C Harrison, Anna A Mikhailova, Elisabeth Marchal, S English, Matthew Carruthers, Emily C Jennings, Emeka L Chiamaka, Ronny A Frigard, Martin Pippel, Geoffrey M Attardo, Joshua B Benoit, Erich Bornberg-Bauer, and Stephen S Tobe. Livebearing cockroach genome reveals convergent evolutionary mechanisms linked to viviparity in insects and beyond. iScience, 26(10):107832, Sep 9 2023. doi: 10.1016/j.isci.2023.107832.

31. Heng Li, Bob Handsaker, Alec Wysoker, Tim Fennell, Jue Ruan, Nils Homer, Gabor Marth, Goncalo Abecasis, and Richard Durbin. The sequence alignment/map format and samtools. Bioinformatics, 25:2078–2079, 8 2009. ISSN 13674803. doi: 10.1093/bioinformatics/btp352.

32. Steven W. Wingett and Simon Andrews. Fastq screen: A tool for multi-genome mapping and quality control [version 1; referees: 3 approved, 1 approved with reservations]. F1000Research, 7, 2018. ISSN 1759796X. doi: 10.12688/f1000research.15931.1.

33. Anthony M. Bolger, Marc Lohse, and Bjoern Usadel. Trimmomatic: A flexible trimmer for illumina sequence data. Bioinformatics, 30:2114–2120, 8 2014. ISSN 14602059. doi: 10.1093/bioinformatics/btu170.

34. Fritz J. Sedlazeck, Philipp Rescheneder, and Arndt Von Haeseler. Nextgenmap: Fast and accurate read mapping in highly polymorphic genomes. Bioinformatics, 29:2790–2791, 11 2013. ISSN 13674803. doi: 10.1093/bioinformatics/btt468.

35. Michael I. Love, Wolfgang Huber, and Simon Anders. Moderated estimation of fold change and dispersion for rna-seq data with deseq2. Genome Biology, 15, 12 2014. ISSN 1474760X. doi: 10.1186/s13059-014-0550-8.

36. Philip Jones, David Binns, Hsin Yu Chang, Matthew Fraser, Weizhong Li, Craig McAnulla, Hamish McWilliam, John Maslen, Alex Mitchell, Gift Nuka, Sebastien Pesseat, Antony F. Quinn, Amaia Sangrador-Vegas, Maxim Scheremetjew, Siew Yit Yong, Rodrigo Lopez, and Sarah Hunter. Interproscan 5: Genome-scale protein function classification. Bioinformatics, 30:1236–1240, 5 2014. ISSN 14602059. doi: 10.1093/bioinformatics/btu031.

37. Adrian Alexa and Jörg Rahnenführer. Gene set enrichment analysis with topgo, 2023.

38. B Bolstad. preprocesscore: A collection of pre-processing functions, 2024.

39. Hadley Wickham. ggplot2: Elegant Graphics for Data Analysis. Springer-Verlag New York, 2016. ISBN 978-3-319-24277-4.

40. David M. Emms and Steven Kelly. Orthofinder: solving fundamental biases in whole genome comparisons dramatically improves orthogroup inference accuracy. Genome Biology, 16, 8 2015. ISSN 1474760X. doi: 10.1186/s13059-015-0721-2.

41. Christiam Camacho, George Coulouris, Vahram Avagyan, Ning Ma, Jason Papadopoulos, Kevin Bealer, and Thomas L. Madden. Blast+: Architecture and applications. BMC Bioinformatics, 10, 12 2009. ISSN 14712105. doi: 10.1186/1471-2105-10-421.

42. JC Oliveros. Venny. an interactive tool for comparing lists with venn’s diagrams., 2007.

43. I Mauger-Birocheau. superexacttest, 2022.

44. Hadley Wickham, Romain François, Lionel Henry, Kirill Müller, and Davis Vaughan. dplyr: A Grammar of Data Manipulation, 2023. R package version 1.1.4, https://github.com/tidyverse/dplyr.

45. Michael Parisi, Rachel Nuttall, Daniel Naiman, Gerard Bouffard, James Malley, Justen Andrews, Scott Eastman, and Brian Oliver. Paucity of genes on the drosophila x chromosome showing male-biased expression. Science, 299(5607):697–700, 2003.

46. José M Ranz, Cristian I Castillo-Davis, Colin D Meiklejohn, and Daniel L Hartl. Sex-dependent gene expression and evolution of the drosophila transcriptome. Science, 300 (5626):1742–1745, 2003.

47. William R Rice. Sex chromosomes and the evolution of sexual dimorphism. Evolution, pages 735–742, 1984.

48. Ann Kathrin Huylmans, Ariana Macon, and Beatriz Vicoso. Global dosage compensation is ubiquitous in lepidoptera, but counteracted by the masculinization of the z chromosome. Molecular Biology and Evolution, 34:2637–2649, 2017. ISSN 15371719. doi: 10.1093/molbev/msx190.

49. Beatriz Vicoso and Doris Bachtrog. Numerous transitions of sex chromosomes in diptera. PLoS biology, 13(4):e1002078, 2015.

50. Ann Kathrin Huylmans and John Parsch. Variation in the x: autosome distribution of male-biased genes among drosophila melanogaster tissues and its relationship with dosage compensation. Genome Biology and Evolution, 7(7):1960–1971, 2015.

51. Liuqi Gu and James R Walters. Evolution of sex chromosome dosage compensation in animals: a beautiful theory, undermined by facts and bedeviled by details. Genome biology and evolution, 9(9):2461–2476, 2017.

52. Peter Luykx. A cytogenetic survey of 25 species of lower termites from australia. Genome, 33(1):80–88, 1990.

53. Francesco Fontana. Interchange complexes in italian populations of reticulitermes lucifugus rossi (isoptera: Rhinotermitidae). Chromosoma, 81(2):169–175, 1980.

54. Kenji Matsuura. A test of the haplodiploid analogy hypothesis in the termite reticulitermes speratus (isoptera: Rhinotermitidae). Annals of the Entomological Society of America, 95 (5):646–649, 2002.

55. Sergio Pimpinelli, M Berloco, Laura Fanti, Patrizio Dimitri, Silvia Bonaccorsi, Enzo Marchetti, Ruggiero Caizzi, C Caggese, and Maurizio Gatti. Transposable elements are stable structural components of drosophila melanogaster heterochromatin. Proceedings of the National Academy of Sciences, 92(9):3804–3808, 1995.

56. Marta Tomaszkiewicz, Paul Medvedev, and Kateryna D Makova. Y and w chromosome assemblies: approaches and discoveries. Trends in genetics, 33(4):266–282, 2017.

57. Sarah B Carey, John T Lovell, Jerry Jenkins, Jim Leebens-Mack, Jeremy Schmutz, Melissa A Wilson, and Alex Harkess. Representing sex chromosomes in genome assemblies. Cell genomics, 2(5), 2022.

58. Karen H Miga, Sergey Koren, Arang Rhie, Mitchell R Vollger, Ariel Gershman, Andrey Bzikadze, Shelise Brooks, Edmund Howe, David Porubsky, Glennis A Logsdon, et al. Telomere-to-telomere assembly of a complete human x chromosome. Nature, 585(7823):79–84, 2020.

59. Lingzhan Xue, Yu Gao, Meiying Wu, Tian Tian, Haiping Fan, Yongji Huang, Zhen Huang, Dapeng Li, and Luohao Xu. Telomere-to-telomere assembly of a fish y chromosome reveals the origin of a young sex chromosome pair. Genome biology, 22:1–20, 2021.

60. Arang Rhie, Sergey Nurk, Monika Cechova, Savannah J Hoyt, Dylan J Taylor, Nicolas Altemose, Paul W Hook, Sergey Koren, Mikko Rautiainen, Ivan A Alexandrov, et al. The complete sequence of a human y chromosome. Nature, 621(7978):344–354, 2023.

61. Kateryna D Makova, Brandon D Pickett, Robert S Harris, Gabrielle A Hartley, Monika Cechova, Karol Pal, Sergey Nurk, DongAhn Yoo, Qiuhui Li, Prajna Hebbar, et al. The complete sequence and comparative analysis of ape sex chromosomes. Nature, pages 1–11, 2024.

62. Doris Bachtrog, Judith E Mank, Catherine L Peichel, Mark Kirkpatrick, Sarah P Otto, Tia-Lynn Ashman, Matthew W Hahn, Jun Kitano, Itay Mayrose, Ray Ming, et al. Sex determination: why so many ways of doing it? PLoS biology, 12(7):e1001899, 2014.

63. Beatriz Vicoso. Molecular and evolutionary dynamics of animal sex-chromosome turnover. Nature ecology & evolution, 3(12):1632–1641, 2019.

64. Lili Wang, Shuran Liao, Minglun Liu, Wenbo Deng, Jiajun He, Zongqing Wang, and Yanli Che. Chromosome number diversity in asian cryptocercus (blattodea, cryptocercidae) and implications for karyotype evolution and geographic distribution on the western sichuan plateau. Systematics and biodiversity, 17(6):594–608, 2019.

65. CA Nalepa, P Luykx, KD Klass, and LL Deitz. Distribution of karyotypes of the cryptocercus punctulatus species complex (dictyoptera: Cryptocercidae) in the southern appalachians: relation to habitat and history. Annals of the Entomological Society of America, 95(3):276– 287, 2002.

66. Ronda L Hamm, Richard P Meisel, and Jeffrey G Scott. The evolving puzzle of autosomal versus y-linked male determination in musca domestica. G3: Genes, Genomes, Genetics, 5(3):371–384, 2015.

67. Krzysztof P Lubieniecki, Song Lin, Emily I Cabana, Jieying Li, Yvonne YY Lai, and William S Davidson. Genomic instability of the sex-determining locus in atlantic salmon (salmo salar). G3: Genes, Genomes, Genetics, 5(11):2513–2522, 2015.

68. Jacob A Tennessen, Na Wei, Shannon CK Straub, Rajanikanth Govindarajulu, Aaron Liston, and Tia-Lynn Ashman. Repeated translocation of a gene cassette drives sex-chromosome turnover in strawberries. PLoS biology, 16(8):e2006062, 2018.

69. Wenlu Yang, Deyan Wang, Yiling Li, Zhiyang Zhang, Shaofei Tong, Mengmeng Li, Xu Zhang, Lei Zhang, Liwen Ren, Xinzhi Ma, et al. A general model to explain repeated turnovers of sex determination in the salicaceae. Molecular Biology and Evolution, 38(3):968–980, 2021.

70. Hans Ellegren and John Parsch. The evolution of sex-biased genes and sex-biased gene expression. Nature Reviews Genetics, 8(9):689–698, 2007.

71. John Parsch and Hans Ellegren. The evolutionary causes and consequences of sex-biased gene expression. Nature Reviews Genetics, 14(2):83–87, 2013.

72. Brian Charlesworth. Model for evolution of y chromosomes and dosage compensation. Proceedings of the National Academy of Sciences, 75(11):5618–5622, 1978.

73. Judith E. Mank, David J. Hosken, and Nina Wedell. Some inconvenient truths about sex chromosome dosage compensation and the potential role of sexual conflict. Evolution, 65 (8):2133–2144, August 2011. doi: 10.1111/j.1558-5646.2011.01316.x.

74. Judith E Mank. Sex chromosome dosage compensation: definitely not for everyone. Trends in genetics, 29(12):677–683, 2013.

