## Supplementary figures and tables for "Recurrent sex chromosome turn-over in termites"

### Supplementary Material for Recurrent sex chromosome turn-over in termites

Roxanne Fraser[1*], Ruth Moraa[1*], Annika Djolai[1], Nils Meisenheimer[1], Sophie Laube[1], Beatriz Vicoso[2], Ann Kathrin Huylmans[1,2✉]
1. Institute of Organismic and Molecular Evolution, Johannes Gutenberg-Universität, Mainz, Germany
2. Institute for Science and Technology Austria, Klosterneuburg, Austria
* These authors contributed equally
✉ Correspondence to: a.huylmans[at]uni-mainz.de

#### 1 Supplementary figures

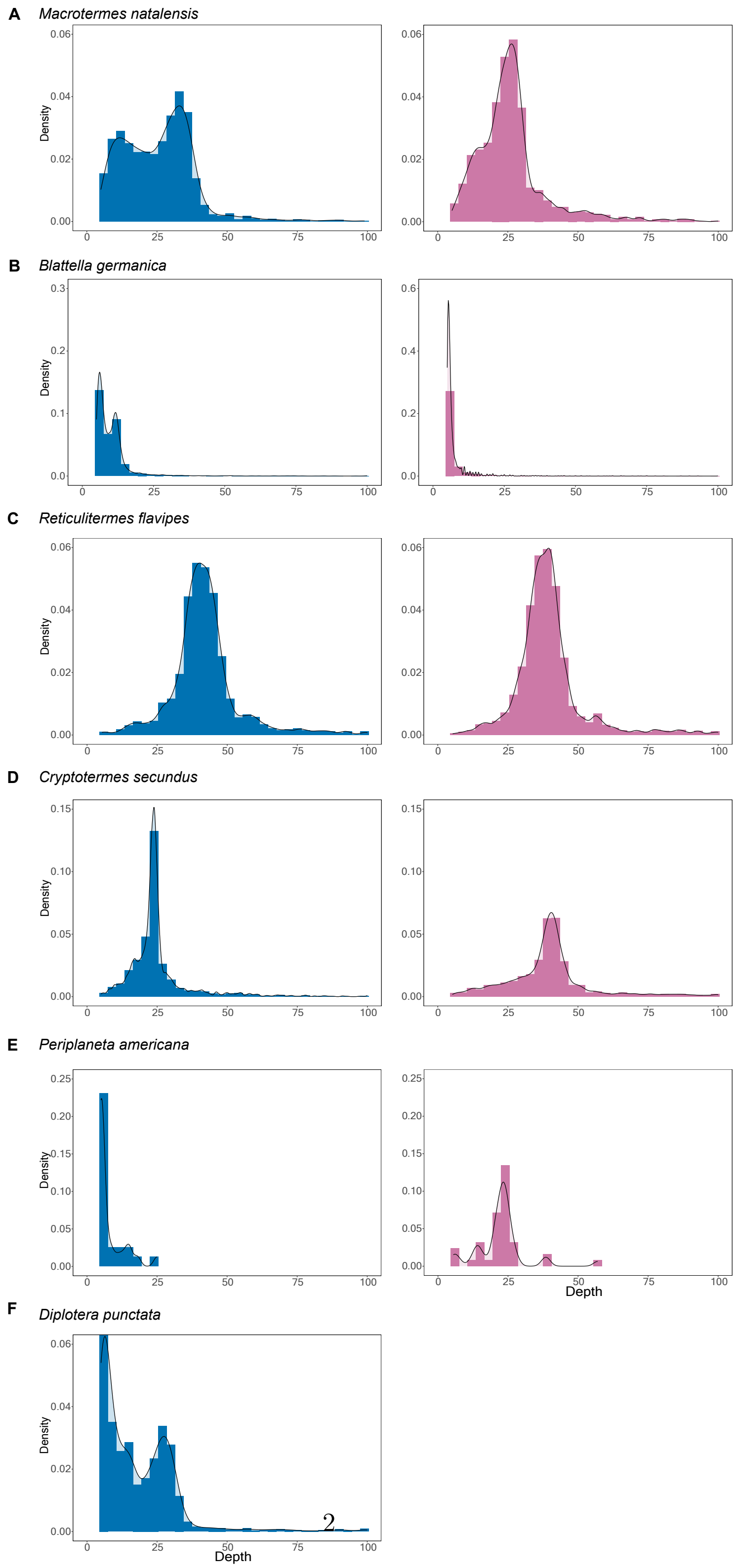


Figure S1: **Male and female genomic coverage of different cockroach and termite species**. Coverage of males (blue) and females (pink) was plotted for (A) *M. natalensis*, (B) *B. germanica*, (C) *R. speratus*, (D) *C. secundus*, and (E) *P. americana*. (F) In *D. punctata*, no female data were available and therefore, only male coverage (blue) was plotted.

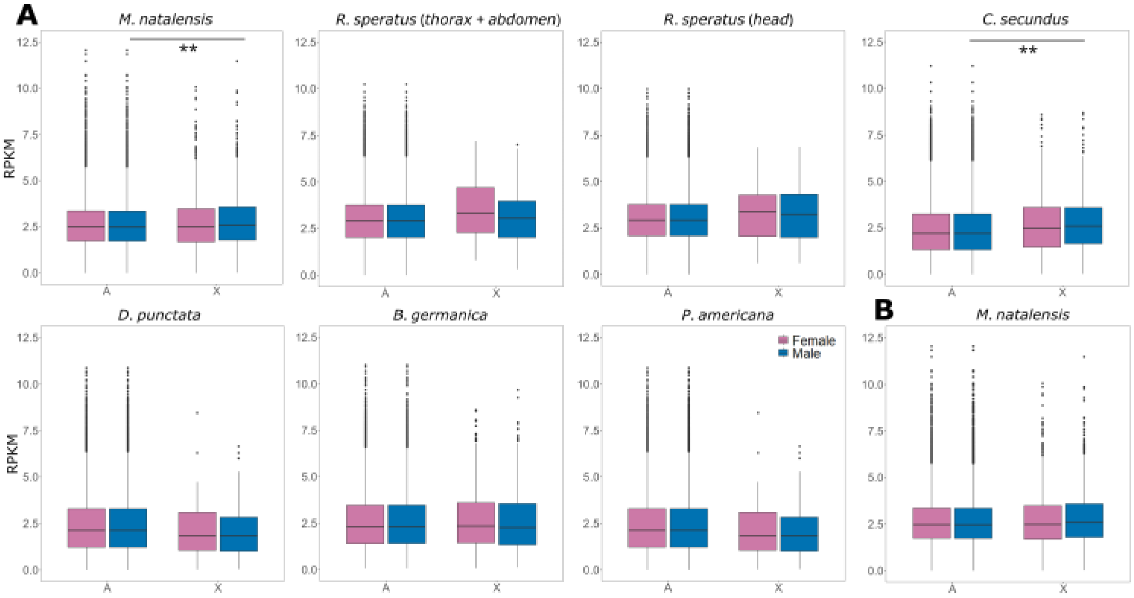


Figure S2: **Dosage compensation adjusts for the chromosomal imbalance in males.** (A) Chromosomal gene expression was compared between males and females for X-linked and autosomal genes in three termite (upper row) and three cockroach species (lower row). For the termite *R. speratus*, no whole body data was available and thus throax and abdomen (including gonads) and head were tested individually. (B) Analysis was repeated after excluding male-biased X-chromosomal genes in *M. natalensis*. Wilcoxon rank sum and signed rank test; * $P < 0.05$; ** $P < 0.01$, *** $P < 0.001$. RPKM = Reads Per Kilobase of transcript per Million mapped reads.

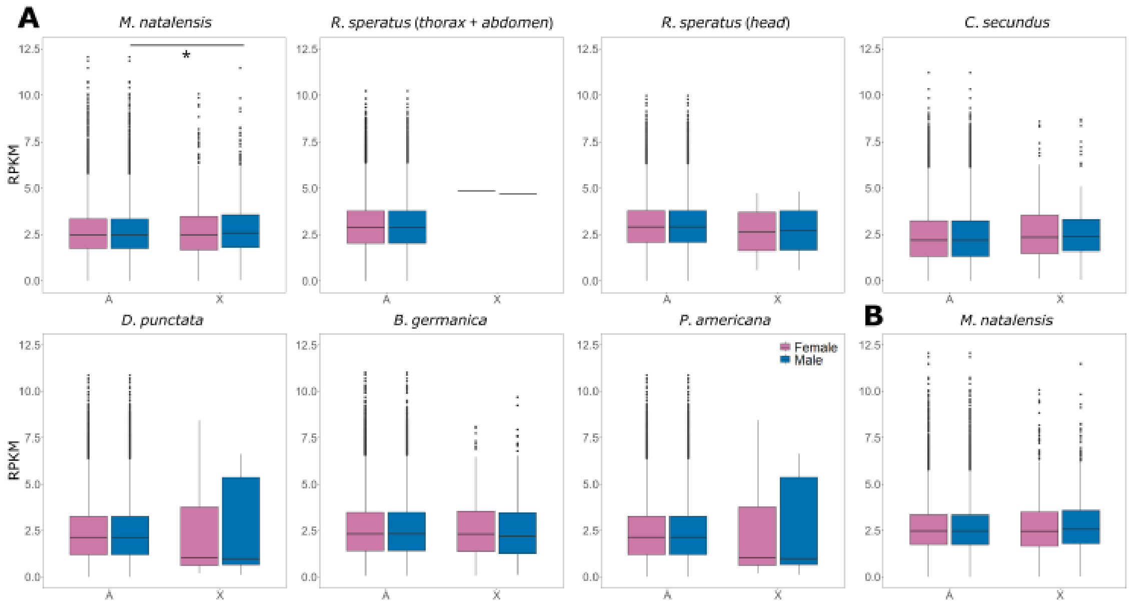


Figure S3: **Dosage compensation also present with more conservative X cutoff.** (A) Chromosomal gene expression was compared between males and females for X-linked and autosomal genes in three termite (upper row) and three cockroach species (lower row), this time for the more conservative X and autosome definition. For the termite *R. speratus*, no whole body data was available and thus throax and abdomen (including gonads) and head were tested individually. (B) In *M. natalensis*, the analysis was repeated after excluding male-biased X-chromosomal genes. Wilcoxon rank sum and signed rank test; * $P < 0.05$; ** $P < 0.01$, *** $P < 0.001$. RPKM = Reads Per Kilobase of transcript per Million mapped reads.

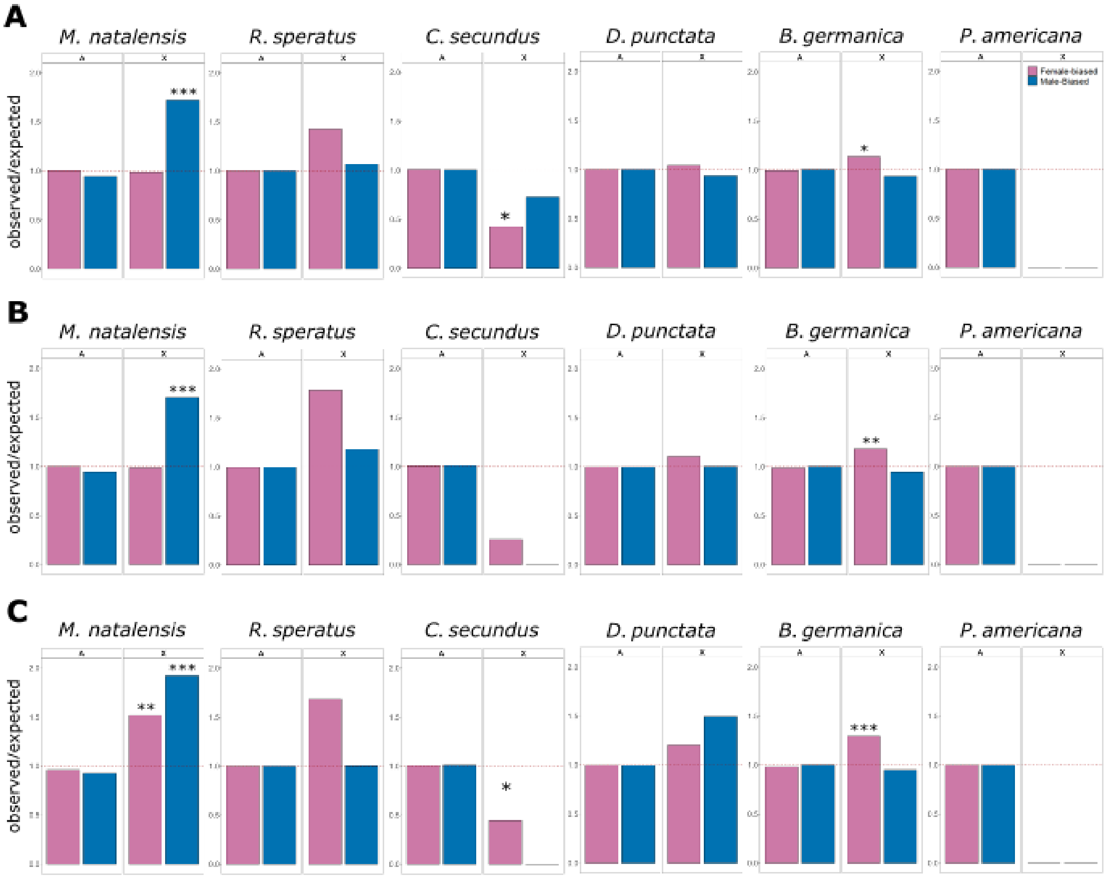


Figure S4: **Enrichment and depletion of sex-biased genes on the X chromosome.** The ratio of observed-to-expected numbers of male- and female-biased genes for (A) all significantly sex-biased genes (no fold-change cutoff), (B) genes with at least medium sex bias (> 2-fold difference between the sexes), and (C) only highly sex-biased genes (> 4-fold difference between the sexes), was plotted for both X chromosome and autosomes. The number of autosomal and X-chromosomal genes was compared between (i) male-biased and unbiased as well as (ii) female-biased and unbiased genes using Fisher's Exact Test; * $P < 0.05$; ** $P < 0.01$, *** $P < 0.001$. Dashed line indicates an observed-to-expected ratio of 1, indicating no deviation from neutral expectations.

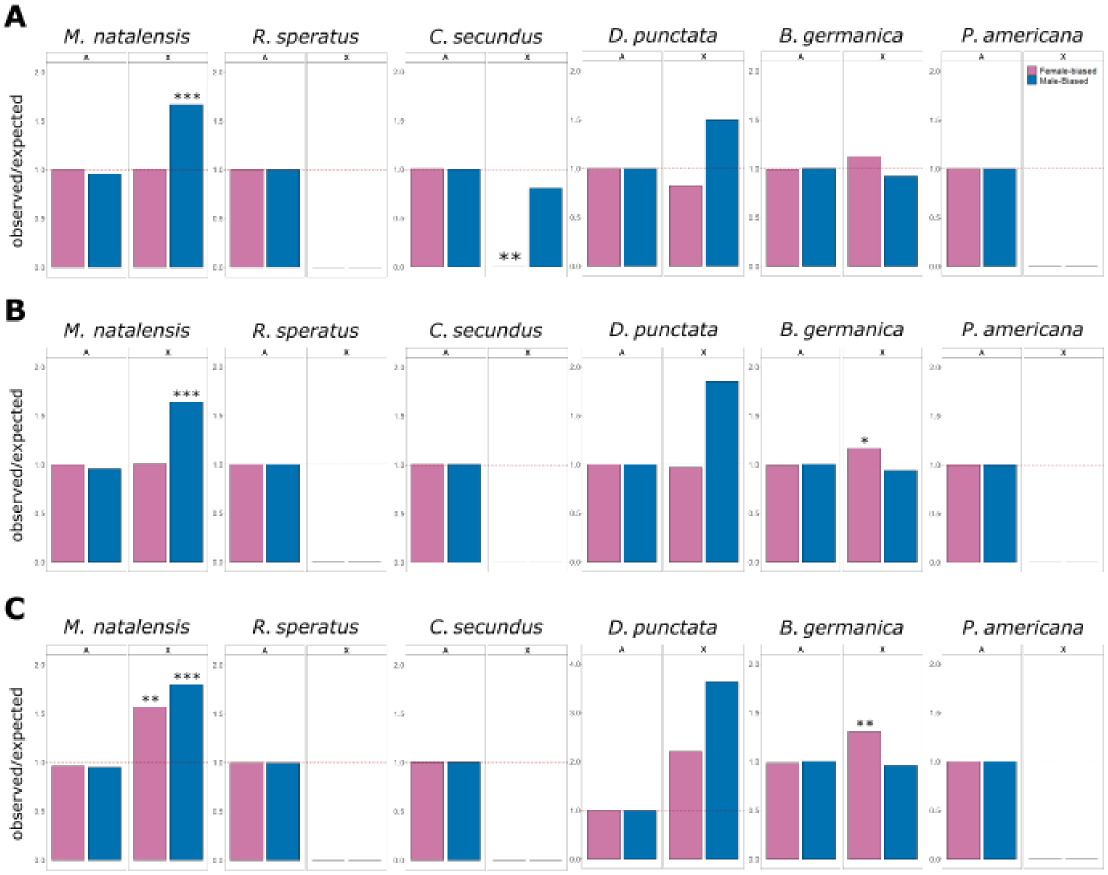


Figure S5: **Enrichment and depletion of sex-biased genes on the X chromosome with more conservative X chromosome cutoff.** The ratio of observed-to-expected numbers of male- and female-biased genes for (A) all significantly sex-biased genes (no fold-change cutoff), (B) genes with at least medium sex bias (> 2-fold difference between the sexes), (C) and only highly sex-biased genes (> 4-fold difference between the sexes) was plotted for both the conservative X chromosome and autosomes. The number of autosomal and X-chromosomal genes was compared between (i) male-biased and unbiased as well as (ii) female-biased and unbiased genes using Fisher's Exact Test; * $P < 0.05$; ** $P < 0.01$, *** $P < 0.001$. Dashed line indicates an observed-to-expected ratio of 1, indicating no deviation from neutral expectations. Please note changed Y axis for highly sex-biased genes in *D. punctata* (C).

#### 2 Supplementary tables

Table S1: **Total number of scaffolds classified as either autosomal (A), X-chromosomal (X), and unknown (NA).**

| Species | A | | X | | NA | | Total |
|---|---|---|---|---|---|---|---|
| ***B. germanica*** | 15,933 | 64.20% | 8,184 | 32.98% | 701 | 2.82% | 24,818 |
| ***P. americana*** | 35 | 72.92% | 13 | 27.08% | 0 | 0.00% | 48 |
| ***D. punctata*** | 9,825 | 91.83% | 869 | 8.12% | 5 | 0.05% | 10,699 |
| ***R. flavipes*** | 5,375 | 92.40% | 409 | 7.03% | 33 | 0.57% | 5,817 |
| ***M. natalensis*** | 29,441 | 20.19% | 41,525 | 28.48% | 74,828 | 51.32% | 145,794 |
| ***C. secundus*** | 73,541 | 25.93% | 50,291 | 17.73% | 159,776 | 56.34% | 283,608 |

Table S2: **Total absolute size of X-chromosomal (X), autosomal (A), and unknown scaffolds (NA) in comparison to the total genome size.**

| | Genome Size | A | | X | | NA | |
|---|---|---|---|---|---|---|---|
| ***B. germanica*** | 2,037,297,555 | 1,787,982,841 | 87.76% | 245,133,537 | 12.03% | 4,181,177 | 0.21% |
| ***P. americana*** | 3,055,948,503 | 3,051,868,968 | 99.87% | 4,079,535 | 0.13% | 0 | 0% |
| ***D. punctata*** | 3,127,968,605 | 3,101,303,600 | 99.15% | 26,635,774 | 0.85% | 29,231 | 0.00% |
| ***R. flavipes*** | 881,338,091 | 878,678,921 | 99.70% | 2,605,511 | 0.3% | 53,659 | 0.00% |
| ***M. natalensis*** | 1,172,292,920 | 1,081,373,577 | 92.24% | 81,414,884 | 6.94% | 9,504,459 | 0.81% |
| ***C. secundus*** | 1,047,420,850 | 1,000,753,519 | 95.54% | 27,330,475 | 2.61% | 19,336,856 | 1.85% |

Table S3: **X-linked genes conserved between *M. natalensis* and *B. germanica*.** Genes which were determined to be conserved between the two species are listed along with their scaffold IDs. Gene pairs marked by an asterisk (*) were also conserved using the more conservative cutoff.

| **Gene Pair** | **Gene ID *M. natalensis*** | **Gene ID *B. germanica*** | **Scaffold ID *M. natalensis*** | **Scaffold ID *B. germanica*** |
|---|---|---|---|---|
| **1*** | Mnat_17507 | Bger_01233 | scaffold9856 | PYGN01000354.1 |
| **2** | Mnat_15962 | Bger_27411 | scaffold1303 | PYGN01003076.1 |
| **3** | Mnat_18012 | Bger_09153 | scaffold1016 | PYGN01000140.1 |
| **4*** | Mnat_17008 | Bger_01236 | scaffold2811 | PYGN01000354.1 |

Table S4: **Gene Ontology terms of conserved genes between *M. natalensis* and *B. germanica***

| Conserved Gene ID | Go Term | Function |
|---|---|---|
| **Mnat_15962** | GO:0006457 | protein folding |
| | GO:0000774 | adenyl-nucleotide exchange factor activity |
| | GO:0042803 | protein homodimerization activity |
| | GO:0051087 | chaperone binding |
| | GO:0001405 | death effector domain binding |
| | GO:0030150 | protein import into mitochondrial matrix |
| | GO:0051082 | unfolded protein binding |
| **Mnat_18012** | GO:0004252 | serine-type endopeptidase activity |
| | GO:0006508 | proteolysis |
| **Bger_27411** | GO:0005887 | Integral component of plasma membrane |
| | GO:0019991 | Monooxygenase activity |
| **Bger_09153** | GO:0004252 | Serine-type endopeptidase activity |
| | GO:0006508 | Proteolysis |

Table S5: **Comparison of numbers of X-linked genes between species pairs.** Fisher's T Test with a P-value $< 0.05$ suggests significant divergence from the random distribution of genes, i.e. more or less X-linked genes shared between two species.

| | **P-value** | |
|---|---|---|
| **Species comparison** | **X chromosome cutoff used in main text** | **More conservative X chromosome cutoff** |
| ***C. secundus* & *B. germanica*** | 0.301 | 1 |
| ***C. secundus* & *R. speratus*** | 1 | 1 |
| ***M. natalensis* & *B. germanica*** | 0.003 | 0.026 |
| ***M. natalensis* & *C. secundus*** | 0.011 | 0.249 |
| ***R. speratus* & *B. germanica*** | 1 | 1 |
| ***R. speratus* & *M. natalensis*** | 0.008 | 1 |
| ***D. punctata* & *B. germanica*** | 0.603 | 1 |
| ***D. punctata* & *C. secundus*** | 0.035 | 0.001 |
| ***D. punctata* & *M. natalensis*** | 0.455 | 1 |
| ***D. punctata* & *P. americana*** | 1 | 1 |
| ***D. punctata* & *R. speratus*** | 1 | 1 |
| ***P. americana* & *B. germanica*** | 0.023 | 1 |
| ***P. americana* & *C. secundus*** | 1 | 1 |
| ***P. americana* & *M. natalensis*** | 1 | 1 |
| ***P. americana* & *R. speratus*** | 1 | 1 |

Table S6: **Observed and expected numbers of X-linked genes in pairwise species comparisons.** Numbers above the diagonal show observed numbers of genes X-linked in both species of the pairwise comparison and numbers below the diagonal show expected numbers to be X-linked based on geneset size and overall X-linked genes as determined via Pearson's Chi-squared Test.

| Species comparison | ***Bger*** | ***Dpun*** | ***Pame*** | ***Mnat*** | ***Csec*** | ***Rspe*** |
|---|---|---|---|---|---|---|
| ***B. germanica (Bger)*** | | 1 | 1 | 4 | 2 | 0 |
| ***D. punctata (Dpun)*** | 0.9002 | | 0 | 1 | 1 | 0 |
| ***P. americana (Pame)*** | 0.0219 | 0.0045 | | 0 | 0 | 0 |
| ***M. natalensis (Mnat)*** | 13.7758 | 0.5949 | 0 | | 4 | 3 |
| ***C. secundus (Csec)*** | 1.0971 | 0.0360 | 0.0669 | 0.9016 | | 0 |
| ***R. speratus (Rspe)*** | 0.641 | 0.0164 | 0.0051 | 0.4351 | 0.0148 | |

Table S7: **Total number of genes showing sex-biased or unbiased expression in termites and cockroaches**. Sex-biased genes are classified according to statistical significance alone without any fold change cut-off. Unbiased genes are those that are expressed but do not show a significant adjusted P-value in the differential expression analysis. Non-expressed genes either had no or very low expression or varied too greatly between biological replicates to calculate any fold change or adjusted P-value.

| Species | Female-biased | Male-biased | Unbiased | Non-expressed | Total |
|---|---|---|---|---|---|
| ***B. germanica*** | 2,650 | 4,914 | 12,735 | 9,814 | 30,113 |
| ***P. americana*** | 308 | 172 | 12,874 | 13,693 | 27,047 |
| ***D. punctata*** | 3,741 | 3,487 | 17,129 | 4,059 | 28,416 |
| ***R. speratus*** | 959 | 1,749 | 11,873 | 1,010 | 15,591 |
| ***M. natalensis*** | 1,202 | 765 | 9,950 | 4,393 | 16,310 |
| ***C. secundus*** | 1,027 | 467 | 13,974 | 2,964 | 18,432 |

Table S8: **Gene ontology terms determined for each species.** For each species, the function of gene ontology term is listed along with the direction of bias, count of significant occurrences, and P-value. Species abbreviation as follows: Bger=*B. germanica*, Pame=*P. americana*, Dpun=*D. punctata* (3 cockroaches), Rspe=*R. speratus*, Mnat=*M. natalensis*, Csec=*C. secundus* (3 termites); FBG=female-biased genes, MBG=male-biased genes.

| GO ID | Term | P-value | Significant | Bias | Species |
|---|---|---|---|---|---|
| **GO:0003674** | molecular_function | 8.50E-20 | 990 | FBG | Bger |
| **GO:0003824** | catalytic activity | 6.50E-18 | 428 | FBG | Bger |
| **GO:0016787** | hydrolase activity | 1.50E-10 | 171 | FBG | Bger |
| **GO:0005576** | extracellular region | 9.60E-08 | 28 | FBG | Bger |
| **GO:0008150** | biological_process | 1.70E-07 | 182 | FBG | Bger |
| **GO:0016491** | oxidoreductase activity | 8.30E-07 | 104 | FBG | Bger |
| **GO:0008152** | metabolic process | 2.80E-06 | 133 | FBG | Bger |
| **GO:0008236** | serine-type peptidase activity | 3.30E-06 | 42 | FBG | Bger |
| **GO:0017171** | serine hydrolase activity | 3.30E-06 | 42 | FBG | Bger |
| **GO:0004497** | monooxygenase activity | 5.90E-06 | 49 | FBG | Bger |
| **GO:0016627** | oxidoreductase activity, acting on the C... | 6.30E-06 | 11 | FBG | Bger |
| **GO:0004601** | peroxidase activity | 1.00E-05 | 13 | FBG | Bger |
| **GO:0016684** | oxidoreductase activity, acting on perox... | 1.00E-05 | 13 | FBG | Bger |
| **GO:0004553** | hydrolase activity, hydrolyzing O-glycos... | 1.10E-05 | 29 | FBG | Bger |
| **GO:0016798** | hydrolase activity, acting on glycosyl b... | 1.10E-05 | 30 | FBG | Bger |
| **GO:0004252** | serine-type endopeptidase activity | 1.50E-05 | 36 | FBG | Bger |
| **GO:1901605** | alpha-amino acid metabolic process | 3.60E-05 | 7 | FBG | Bger |
| **GO:0004096** | catalase activity | 5.90E-05 | 10 | FBG | Bger |
| **GO:0003924** | GTPase activity | 8.90E-05 | 36 | FBG | Bger |
| **GO:0016209** | antioxidant activity | 0.00011 | 13 | FBG | Bger |
| **GO:0071704** | organic substance metabolic process | 0.00011 | 118 | FBG | Bger |
| **GO:0015078** | proton transmembrane transporter activit... | 0.00012 | 12 | FBG | Bger |
| **GO:0006790** | sulfur compound metabolic process | 0.00014 | 7 | FBG | Bger |
| **GO:0000096** | sulfur amino acid metabolic process | 0.00018 | 6 | FBG | Bger |
| **GO:0006534** | cysteine metabolic process | 0.00018 | 6 | FBG | Bger |
| **GO:0009069** | serine family amino acid metabolic proce... | 0.00018 | 6 | FBG | Bger |
| **GO:0009092** | homoserine metabolic process | 0.00018 | 6 | FBG | Bger |
| **GO:0019346** | transsulfuration | 0.00018 | 6 | FBG | Bger |
| **O:0050667** | homocysteine metabolic process | 0.00018 | 6 | FBG | Bger |
| **GO:0017111** | ribonucleoside triphosphate phosphatase ... | 0.00019 | 36 | FBG | Bger |
| **GO:0005975** | carbohydrate metabolic process | 0.00021 | 25 | FBG | Bger |
| **GO:0044238** | primary metabolic process | 0.00028 | 115 | FBG | Bger |
| **GO:0003674** | molecular_function | 3.00E-22 | 1104 | MBG | Bger |
| **GO:0035639** | purine ribonucleoside triphosphate bindi... | 2.90E-12 | 86 | MBG | Bger |
| **GO:0017076** | purine nucleotide binding | 7.50E-12 | 87 | MBG | Bger |
| **GO:0032555** | purine ribonucleotide binding | 7.50E-12 | 87 | MBG | Bger |
| **GO:0032553** | ribonucleotide binding | 1.30E-11 | 87 | MBG | Bger |
| **GO:0097367** | carbohydrate derivative binding | 2.20E-11 | 87 | MBG | Bger |
| **GO:0043168** | anion binding | 6.30E-11 | 87 | MBG | Bger |

| | | | | | |
|---|---|---|---|---|---|
| **GO:0005524** | ATP binding | 3.90E-09 | 68 | MBG | Bger |
| **GO:0016020** | membrane | 4.00E-09 | 262 | MBG | Bger |
| **GO:0030554** | adenyl nucleotide binding | 9.00E-09 | 69 | MBG | Bger |
| **GO:0032559** | adenyl ribonucleotide binding | 9.00E-09 | 69 | MBG | Bger |
| **GO:0000166** | nucleotide binding | 2.30E-08 | 92 | MBG | Bger |
| **GO:1901265** | nucleoside phosphate binding | 2.30E-08 | 92 | MBG | Bger |
| **GO:0036094** | small molecule binding | 4.90E-08 | 92 | MBG | Bger |
| **GO:0022836** | gated channel activity | 1.70E-07 | 94 | MBG | Bger |
| **GO:0015276** | ligand-gated ion channel activity | 1.70E-07 | 93 | MBG | Bger |
| **GO:0022834** | ligand-gated channel activity | 1.70E-07 | 93 | MBG | Bger |
| **GO:0043167** | ion binding | 6.80E-07 | 130 | MBG | Bger |
| **GO:0008150** | biological_process | 1.10E-06 | 162 | MBG | Bger |
| **GO:0097159** | organic cyclic compound binding | 1.80E-06 | 239 | MBG | Bger |
| **GO:1901363** | heterocyclic compound binding | 1.80E-06 | 239 | MBG | Bger |
| **GO:0005216** | ion channel activity | 4.60E-06 | 115 | MBG | Bger |
| **GO:0015267** | channel activity | 4.90E-06 | 116 | MBG | Bger |
| **GO:0022803** | passive transmembrane transporter activi... | 4.90E-06 | 116 | MBG | Bger |
| **GO:0015318** | inorganic molecular entity transmembrane... | 6.20E-06 | 121 | MBG | Bger |
| **GO:0015075** | ion transmembrane transporter activity | 9.30E-06 | 120 | MBG | Bger |
| **GO:0110165** | cellular anatomical entity | 2.00E-05 | 314 | MBG | Bger |
| **GO:0004185** | serine-type carboxypeptidase activity | 9.50E-05 | 11 | MBG | Bger |
| **GO:0070008** | serine-type exopeptidase activity | 9.50E-05 | 11 | MBG | Bger |
| **GO:0022857** | transmembrane transporter activity | 0.00013 | 138 | MBG | Bger |
| **GO:0005215** | transporter activity | 0.00013 | 140 | MBG | Bger |
| **GO:0005525** | GTP binding | 0.00015 | 18 | MBG | Bger |
| **GO:0019001** | guanyl nucleotide binding | 0.00015 | 18 | MBG | Bger |
| **GO:0032561** | guanyl ribonucleotide binding | 0.00015 | 18 | MBG | Bger |
| **GO:0003824** | catalytic activity | 4.10E-11 | 60 | FBG | Pame |
| **GO:0008236** | serine-type peptidase activity | 2.10E-09 | 13 | FBG | Pame |
| **GO:0017171** | serine hydrolase activity | 2.10E-09 | 13 | FBG | Pame |
| **GO:0016491** | oxidoreductase activity | 2.50E-09 | 23 | FBG | Pame |
| **GO:0004252** | serine-type endopeptidase activity | 6.90E-09 | 12 | FBG | Pame |
| **GO:0016798** | hydrolase activity, acting on glycosyl b... | 1.80E-08 | 11 | FBG | Pame |
| **GO:0005506** | iron ion binding | 2.50E-08 | 14 | FBG | Pame |
| **GO:0004553** | hydrolase activity, hydrolyzing O-glycos... | 8.00E-08 | 10 | FBG | Pame |
| **GO:0020037** | heme binding | 9.10E-08 | 14 | FBG | Pame |
| **GO:0046906** | tetrapyrrole binding | 9.10E-08 | 14 | FBG | Pame |
| **GO:0004497** | monooxygenase activity | 2.80E-07 | 13 | FBG | Pame |
| **GO:0016705** | oxidoreductase activity, acting on paire... | 2.80E-07 | 13 | FBG | Pame |
| **GO:0016787** | hydrolase activity | 4.00E-07 | 32 | FBG | Pame |
| **GO:0004175** | endopeptidase activity | 1.00E-06 | 12 | FBG | Pame |
| **GO:0006508** | proteolysis | 2.10E-06 | 16 | FBG | Pame |
| **GO:0008233** | peptidase activity | 2.20E-06 | 15 | FBG | Pame |
| **GO:0005975** | carbohydrate metabolic process | 3.40E-06 | 12 | FBG | Pame |
| **GO:0046914** | transition metal ion binding | 1.00E-05 | 17 | FBG | Pame |
| **GO:0046872** | metal ion binding | 5.80E-05 | 19 | FBG | Pame |

| | | | | | |
|---|---|---|---|---|---|
| **GO:0043169** | cation binding | 6.90E-05 | 19 | FBG | Pame |
| **GO:0000786** | nucleosome | 6.90E-05 | 3 | MBG | Pame |
| **GO:0000785** | chromatin | 0.00011 | 3 | MBG | Pame |
| **GO:0032993** | protein-DNA complex | 0.00011 | 3 | MBG | Pame |
| **GO:0044815** | DNA packaging complex | 0.00011 | 3 | MBG | Pame |
| **GO:0015276** | ligand-gated ion channel activity | 1.20E-19 | 81 | FBG | Dpun |
| **GO:0022834** | ligand-gated channel activity | 1.20E-19 | 81 | FBG | Dpun |
| **GO:0003674** | molecular_function | 1.30E-29 | 630 | FBG | Dpun |
| **GO:0005216** | ion channel activity | 1.80E-19 | 99 | FBG | Dpun |
| **GO:0022857** | transmembrane transporter activity | 2.60E-15 | 111 | FBG | Dpun |
| **GO:0022836** | gated channel activity | 3.00E-19 | 81 | FBG | Dpun |
| **GO:0005575** | cellular_component | 5.30E-13 | 230 | FBG | Dpun |
| **GO:0015267** | channel activity | 6.40E-19 | 99 | FBG | Dpun |
| **GO:0022803** | passive transmembrane transporter activi... | 6.40E-19 | 99 | FBG | Dpun |
| **GO:0015075** | ion transmembrane transporter activity | 6.50E-18 | 100 | FBG | Dpun |
| **GO:0110165** | cellular anatomical entity | 7.60E-13 | 227 | FBG | Dpun |
| **GO:0015318** | inorganic molecular entity transmembrane... | 7.70E-18 | 100 | FBG | Dpun |
| **GO:0016020** | membrane | 9.60E-10 | 180 | FBG | Dpun |
| **GO:0005215** | transporter activity | 9.00E-15 | 112 | FBG | Dpun |
| **GO:0043229** | intracellular organelle | 0.00014 | 58 | MBG | Dpun |
| **GO:0043226** | organelle | 0.00021 | 58 | MBG | Dpun |
| **GO:0005575** | cellular_component | 1.10E-09 | 172 | MBG | Dpun |
| **GO:0140096** | catalytic activity, acting on a protein | 1.70E-05 | 139 | MBG | Dpun |
| **GO:0008233** | peptidase activity | 1.90E-07 | 89 | MBG | Dpun |
| **GO:0008150** | biological_process | 1.90E-08 | 206 | MBG | Dpun |
| **GO:0004252** | serine-type endopeptidase activity | 2.00E-06 | 55 | MBG | Dpun |
| **GO:0016491** | oxidoreductase activity | 2.10E-11 | 111 | MBG | Dpun |
| **GO:0008236** | serine-type peptidase activity | 2.30E-06 | 58 | MBG | Dpun |
| **GO:0017171** | serine hydrolase activity | 2.30E-06 | 58 | MBG | Dpun |
| **GO:0003824** | catalytic activity | 2.60E-19 | 427 | MBG | Dpun |
| **GO:0016787** | hydrolase activity | 2.80E-11 | 190 | MBG | Dpun |
| **GO:0004497** | monooxygenase activity | 5.80E-07 | 49 | MBG | Dpun |
| **GO:0005622** | intracellular anatomical structure | 6.20E-05 | 64 | MBG | Dpun |
| **GO:0005635** | nuclear envelope | 7.20E-05 | 6 | MBG | Dpun |
| **GO:0005643** | nuclear pore | 7.20E-05 | 6 | MBG | Dpun |
| **GO:0004175** | endopeptidase activity | 7.50E-07 | 68 | MBG | Dpun |
| **GO:0110165** | cellular anatomical entity | 8.80E-06 | 165 | MBG | Dpun |
| **GO:0003674** | molecular_function | 1.70E-18 | 316 | FBG | Rspe |
| **GO:0015267** | channel activity | 4.30E-05 | 22 | FBG | Rspe |
| **GO:0022803** | passive transmembrane transporter activi... | 4.30E-05 | 22 | FBG | Rspe |
| **GO:0005549** | odorant binding | 5.00E-28 | 24 | FBG | Rspe |
| **GO:0008150** | biological_process | 1.20E-07 | 112 | MBG | Rspe |
| **GO:0008233** | peptidase activity | 1.20E-08 | 57 | MBG | Rspe |
| **GO:0071704** | organic substance metabolic process | 1.40E-05 | 76 | MBG | Rspe |
| **GO:0110165** | cellular anatomical entity | 1.70E-08 | 126 | MBG | Rspe |
| **GO:0004175** | endopeptidase activity | 2.30E-07 | 42 | MBG | Rspe |

| | | | | | |
|---|---|---|---|---|---|
| **GO:0016787** | hydrolase activity | 2.70E-12 | 128 | MBG | Rspe |
| **GO:0003824** | catalytic activity | 3.90E-13 | 276 | MBG | Rspe |
| **GO:0044238** | primary metabolic process | 4.10E-06 | 76 | MBG | Rspe |
| **GO:0004497** | monooxygenase activity | 4.10E-07 | 36 | MBG | Rspe |
| **GO:0016491** | oxidoreductase activity | 4.30E-08 | 68 | MBG | Rspe |
| **GO:0008236** | serine-type peptidase activity | 4.80E-07 | 31 | MBG | Rspe |
| **GO:0017171** | serine hydrolase activity | 4.80E-07 | 31 | MBG | Rspe |
| **GO:0005975** | carbohydrate metabolic process | 5.20E-06 | 18 | MBG | Rspe |
| **GO:0008152** | metabolic process | 5.90E-08 | 87 | MBG | Rspe |
| **GO:0005575** | cellular_component | 8.40E-08 | 127 | MBG | Rspe |
| **GO:0004252** | serine-type endopeptidase activity | 9.70E-07 | 30 | MBG | Rspe |
| **GO:0034645** | cellular macromolecule biosynthetic proc... | 7.30E-18 | 59 | FBG | Mnat |
| **GO:1901566** | organonitrogen compound biosynthetic pro... | 2.00E-17 | 69 | FBG | Mnat |
| **GO:0006518** | peptide metabolic process | 2.40E-16 | 52 | FBG | Mnat |
| **GO:0003735** | structural constituent of ribosome | 3.80E-16 | 37 | FBG | Mnat |
| **GO:0043603** | cellular amide metabolic process | 6.50E-16 | 52 | FBG | Mnat |
| **GO:0006412** | translation | 9.50E-16 | 49 | FBG | Mnat |
| **GO:0043043** | peptide biosynthetic process | 1.80E-15 | 49 | FBG | Mnat |
| **GO:0043604** | amide biosynthetic process | 1.80E-15 | 49 | FBG | Mnat |
| **GO:0005198** | structural molecule activity | 7.90E-15 | 39 | FBG | Mnat |
| **GO:0005840** | ribosome | 1.60E-11 | 36 | FBG | Mnat |
| **GO:0009059** | macromolecule biosynthetic process | 1.20E-10 | 82 | FBG | Mnat |
| **GO:0005622** | intracellular anatomical structure | 1.80E-10 | 111 | FBG | Mnat |
| **GO:0044249** | cellular biosynthetic process | 2.60E-10 | 94 | FBG | Mnat |
| **GO:1901576** | organic substance biosynthetic process | 5.00E-10 | 95 | FBG | Mnat |
| **GO:0044260** | cellular macromolecule metabolic process | 6.30E-10 | 73 | FBG | Mnat |
| **GO:0043226** | organelle | 1.10E-09 | 97 | FBG | Mnat |
| **GO:0043229** | intracellular organelle | 2.10E-09 | 96 | FBG | Mnat |
| **GO:0044271** | cellular nitrogen compound biosynthetic ... | 2.40E-09 | 83 | FBG | Mnat |
| **GO:0009058** | biosynthetic process | 2.40E-09 | 97 | FBG | Mnat |
| **GO:0010467** | gene expression | 3.60E-09 | 90 | FBG | Mnat |
| **GO:0043228** | non-membrane-bounded organelle | 4.50E-08 | 48 | FBG | Mnat |
| **GO:0043232** | intracellular non-membrane-bounded organ... | 4.50E-08 | 48 | FBG | Mnat |
| **GO:0034641** | cellular nitrogen compound metabolic pro... | 4.90E-07 | 111 | FBG | Mnat |
| **GO:0015078** | proton transmembrane transporter activit... | 4.10E-06 | 15 | FBG | Mnat |
| **GO:0098796** | membrane protein complex | 4.10E-06 | 25 | FBG | Mnat |
| **GO:0009055** | electron transfer activity | 6.00E-06 | 11 | FBG | Mnat |
| **GO:0045263** | proton-transporting ATP synthase complex... | 9.20E-06 | 8 | FBG | Mnat |
| **GO:0045259** | proton-transporting ATP synthase complex | 1.10E-05 | 9 | FBG | Mnat |
| **GO:1901564** | organonitrogen compound metabolic proces... | 2.80E-05 | 116 | FBG | Mnat |
| **GO:0044237** | cellular metabolic process | 3.40E-05 | 150 | FBG | Mnat |
| **GO:0032991** | protein-containing complex | 4.00E-05 | 59 | FBG | Mnat |
| **GO:1901137** | carbohydrate derivative biosynthetic pro... | 4.80E-05 | 19 | FBG | Mnat |
| **GO:0006754** | ATP biosynthetic process | 0.00016 | 8 | FBG | Mnat |
| **GO:0009142** | nucleoside triphosphate biosynthetic pro... | 0.00016 | 8 | FBG | Mnat |
| **GO:0009145** | purine nucleoside triphosphate biosynthe... | 0.00016 | 8 | FBG | Mnat |

| | | | | | |
|---|---|---|---|---|---|
| **GO:0009201** | ribonucleoside triphosphate biosynthetic... | 0.00016 | 8 | FBG | Mnat |
| **GO:0009206** | purine ribonucleoside triphosphate biosy... | 0.00016 | 8 | FBG | Mnat |
| **GO:0015986** | proton motive force-driven ATP synthesis | 0.00016 | 8 | FBG | Mnat |
| **GO:0019538** | protein metabolic process | 0.00018 | 101 | FBG | Mnat |
| **GO:0006807** | nitrogen compound metabolic process | 0.00034 | 163 | FBG | Mnat |
| **GO:0033177** | proton-transporting two-sector ATPase co... | 4.00E-04 | 8 | FBG | Mnat |
| **GO:0031090** | organelle membrane | 0.00042 | 22 | FBG | Mnat |
| **GO:0043170** | macromolecule metabolic process | 0.00046 | 150 | FBG | Mnat |
| **GO:0005634** | nucleus | 5.90E-08 | 31 | FBG | Csec |
| **GO:0005622** | intracellular anatomical structure | 8.00E-06 | 51 | FBG | Csec |
| **GO:0043231** | intracellular membrane-bounded organelle | 8.20E-05 | 33 | FBG | Csec |
| **GO:0043227** | membrane-bounded organelle | 1.00E-04 | 33 | FBG | Csec |
| **GO:0043229** | intracellular organelle | 0.00013 | 42 | FBG | Csec |
| **GO:0043226** | organelle | 0.00015 | 42 | FBG | Csec |
| **GO:0016491** | oxidoreductase activity | 1.00E-12 | 35 | MBG | Csec |
| **GO:0004497** | monooxygenase activity | 4.90E-08 | 15 | MBG | Csec |
| **GO:0016705** | oxidoreductase activity, acting on paire... | 5.60E-08 | 15 | MBG | Csec |
| **GO:0020037** | heme binding | 5.60E-07 | 14 | MBG | Csec |
| **GO:0046906** | tetrapyrrole binding | 6.30E-07 | 14 | MBG | Csec |
| **GO:0005506** | iron ion binding | 2.70E-06 | 13 | MBG | Csec |
| **GO:0046872** | metal ion binding | 7.30E-05 | 33 | MBG | Csec |
| **GO:0043169** | cation binding | 9.20E-05 | 33 | MBG | Csec |

Table S9: **Gene ontology terms enriched in sex-biased genes and conserved within roaches.** FBG: female-biased genes, MBG: male-biased genes.

| **GO.ID** | **Term.x** | ***B. germanica*** | ***P. americana*** | ***D. punctata*** |
|---|---|---|---|---|
| **GO:0003824** | catalytic activity | FBG | FBG | MBG |
| **GO:0016787** | hydrolase activity | FBG | FBG | MBG |
| **GO:0016491** | oxidoreductase activity | FBG | FBG | MBG |
| **GO:0008236** | serine-type peptidase activity | FBG | FBG | MBG |
| **GO:0017171** | serine hydrolase activity | FBG | FBG | MBG |
| **GO:0004497** | monooxygenase activity | FBG | FBG | MBG |
| **GO:0004252** | serine-type endopeptidase activity | FBG | FBG | MBG |

Table S10: **Number of sex-biased and unbiased genes on the X chromosome and autosomes, independently of their fold change.**

| Chromosome | X chromosome | | | Autosome | | |
|---|---|---|---|---|---|---|
| Sex bias | Female-biased | Male-biased | Unbiased | Female-biased | Male-biased | Unbiased |
| ***B. germanica*** | 254 | 256 | 745 | 3654 | 4844 | 12803 |
| ***P. americana*** | 0 | 0 | 6 | 308 | 172 | 12868 |
| ***D. punctata*** | 16 | 13 | 60 | 3655 | 3409 | 14505 |
| ***R. speratus*** | 3 | 4 | 24 | 909 | 1442 | 11475 |
| ***M. natalensis*** | 83 | 98 | 699 | 1119 | 667 | 9251 |
| ***C. secundus*** | 5 | 4 | 172 | 1010 | 454 | 14043 |

Table S11: **Number of sex-biased and unbiased genes on the X chromosome and autosomes with more than 2-fold difference between the sexes.**

| Chromosome | X chromosome | | | Autosome | | |
|---|---|---|---|---|---|---|
| Sex bias | Female-biased | Male-biased | Unbiased | Female-biased | Male-biased | Unbiased |
| ***B. germanica*** | 245 | 245 | 745 | 3339 | 4564 | 12803 |
| ***P. americana*** | 0 | 0 | 6 | 308 | 172 | 12868 |
| ***D. punctata*** | 14 | 11 | 60 | 2994 | 2647 | 14505 |
| ***R. speratus*** | 3 | 4 | 24 | 705 | 1259 | 11475 |
| ***M. natalensis*** | 82 | 96 | 699 | 1101 | 662 | 9251 |
| ***C. secundus*** | 2 | 0 | 172 | 661 | 218 | 14043 |

Table S12: **Number of sex-biased and unbiased genes on the X chromosome and autosomes with more than 4-fold difference between the sexes.**

| **Chromosome** | **X chromosome** | | | **Autosome** | | |
|---|---|---|---|---|---|---|
| **Sex bias** | **Female-biased** | **Male-biased** | **Unbiased** | **Female-biased** | **Male-biased** | **Unbiased** |
| ***B. germanica*** | 157 | 174 | 745 | 1944 | 3196 | 12803 |
| ***P. americana*** | 0 | 0 | 6 | 308 | 168 | 12868 |
| ***D. punctata*** | 6 | 8 | 60 | 1184 | 1236 | 14505 |
| ***R. speratus*** | 1 | 2 | 24 | 233 | 639 | 11475 |
| ***M. natalensis*** | 45 | 65 | 699 | 368 | 396 | 9251 |
| ***C. secundus*** | 1 | 0 | 172 | 187 | 42 | 14043 |
